# Ecological diversification in sexual and asexual lineages

**DOI:** 10.1101/2024.03.06.583698

**Authors:** P. Catalina Chaparro-Pedraza, Gregory Roth, Carlos J. Melián

## Abstract

The presence or absence of sex can have a strong influence on the processes whereby species arise. Yet, the mechanistic underpinnings of this influence are poorly understood. To gain insights into the mechanisms whereby the reproductive mode may influence diversification, we investigate how natural selection, genetic mixing and the reproductive mode interact and how this interaction affects the evolutionary dynamics of diversifying lineages. To do so, we formulate and analyze trait-based eco-evolutionary models of ecological diversification for sexual and asexual lineages, in which diversification is driven by intraspecific resource competition. We find that the reproductive mode strongly influences the diversification rate and thus the ensuing diversity of a lineage. Our results reveal that natural selection is stronger in asexual lineages because asexual organisms have a higher reproductive potential than sexual ones. As a consequence, an asexual population can reach a higher population density than a sexual population under the very same ecological conditions. This causes competition, and thus ecologically-based selection, to be stronger in asexual lineages, promoting faster diversification. However, a small amount of genetic mixing accelerates the trait expansion process in sexual lineages, overturning the effect of selection alone and enabling a faster niche occupancy than asexual lineages. As a consequence, sexual lineages can occupy more ecological niches, eventually resulting in higher diversity. This suggests that sexual reproduction may be widespread among species because it increases rates of diversification.

## Introduction

Ecological diversification, the process whereby new species emerge as a consequence of ecologically-based disruptive selection^1^, is thought to have given rise to much of the diversity of life on Earth ^2,3^. This process has produced several diverse adaptive radiations including Darwin’s finches ^4^, Caribbean anole lizards ^5^, and cichlid fishes ^6^. To understand the origin of biodiversity, it is therefore fundamental to gain insights into the factors influencing ecological diversification. One of these factors may be the presence or absence of sex ^7^.

A wide variety of ecological interactions can induce selection regimes leading to ecological diversification ^8,9^. Arguably, the most studied is intraspecific competition, which can induce disruptive selection in the presence of ecological opportunity ^10,11^, i.e. the availability of relatively unexploited ecological niches ^12–14^. Under disruptive selection, intermediate phenotypes have a fitness disadvantage compared with more extreme phenotypes, causing phenotypic diversification ^13,15,16^. This results in speciation in asexual populations due to the lack of genetic mixing. However, in sexual populations, for speciation to occur, barriers to gene flow must evolve between clusters of individuals with divergent phenotypes ^1,13,17^.

The reproductive mode can additionally influence ecological diversification in multiple manners: On one hand, sexual reproduction, by reshuffling the genetic material, can increase rates of adaptation and evolution ^18–21^. Genetic mixing may perhaps result in higher rates of diversification. On the other hand, an asexual population will have twice the reproductive potential of a sexual population (i.e. the twofold cost of sex due to the production of males, as outlined by Maynard-Smith) ^22^ and therefore can quickly reach a higher population density. With higher population density, competition for resources can be stronger in an asexual population, resulting in stronger ecologically-based disruptive selection driving diversification. Hence, to understand how the reproductive mode influences ecological diversification, it is fundamental to gain insights into the interaction between the reproductive mode, natural selection, and genetic mixing.

Previous research examining the effect of the reproductive mode on diversification has not investigated this interaction. For example, Melián et al. (2012) examined how the reproductive mode influences diversification in a model of neutral evolutionary change, thus neglecting natural selection. Other studies have attempted to explain how clusters may emerge under alternative reproductive modes neglecting the differences in reproductive potential inherent to the reproductive modes^24–26^ (i.e. the twofold cost of sex), which may alter the strength of natural selection and thus diversification. Theory that mechanistically link ecological and microevolutionary processes with diversification patterns is needed to understand how the interaction between the reproductive mode, natural selection and genetic mixing influences ecological diversification.

Here we address this question by analyzing models of ecological diversification driven by intraspecific resource competition. Following the large body of theory in adaptive dynamics ^8,12,13,27^, we model the eco-evolutionary dynamics of a diversifying lineage, which initiates with a founder population colonizing an environment with a variety of unexploited food resources. Through an adaptive process, the lineage diversifies to occupy the available niches. We use a model that was formulated for asexual lineages^28,29^, and a modified variant for sexual lineages to compare the diversification process between the lineages with the alternative reproductive modes, and examine the effect of natural selection and genetic mixing on their evolutionary dynamics. The only ecological difference between the sexual and asexual lineage lies on the fact that asexual females produce only females (i.e., all individuals can produce offspring), whereas sexual females produce females and males with equal probability (i.e. primary sex ratio is 1:1, and only females can produce offspring). Therefore, sexual and asexual females produce the same number of total offspring if they have identical feeding niche trait and experience the same environment. However, because asexuals do not produce male offspring, they produce exactly twice as many female offspring as the sexuals. In some organisms, the cost of producing males need not be this high, e.g., if the sex ratio is biased towards females or if males provide parental care ^30^. Our assumption is therefore one of the least favorable for the sexual populations.

## Methods

We formulate diversification models for sexual and asexual lineages that differ only in the mode of reproduction, whereas environmental conditions (i.e. food resource availability) and ecological demographic rates (i.e. resource use, mortality) are set equal. Then, we use analytical techniques, numerical simulations and individual-based models to examine the effect of natural selection and genetic mixing on diversification.

### Model assumptions

#### The environment: Food resources

We consider *n* food resources to exist prior to arrival of the diversifying lineage (preexisting resources, *i =* 1, …, *n*) with density *R*_i_. In the absence of consumers, resource density dynamics follow 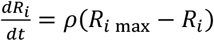, where *ρ* and *R*_*i* max_ are the renewal rate and the carrying capacity of the *i*th resource, respectively. The total productivity is the sum of the carrying capacities of the resources, *P =* ∑_*i*_ *F*_*i max*_. We assume that, for each resource, there exists an optimal trait value *θ*_*i*_ to consume it. For the sake of simplicity, these optimal traits are assumed to be ordered along a one-dimensional ecological trait space (i.e. *θ*_1_ < *θ*_2_ < … < *θ*_*n*_) and equally distant from one another by a distance *D* (figure 1).

**Figure 1.**
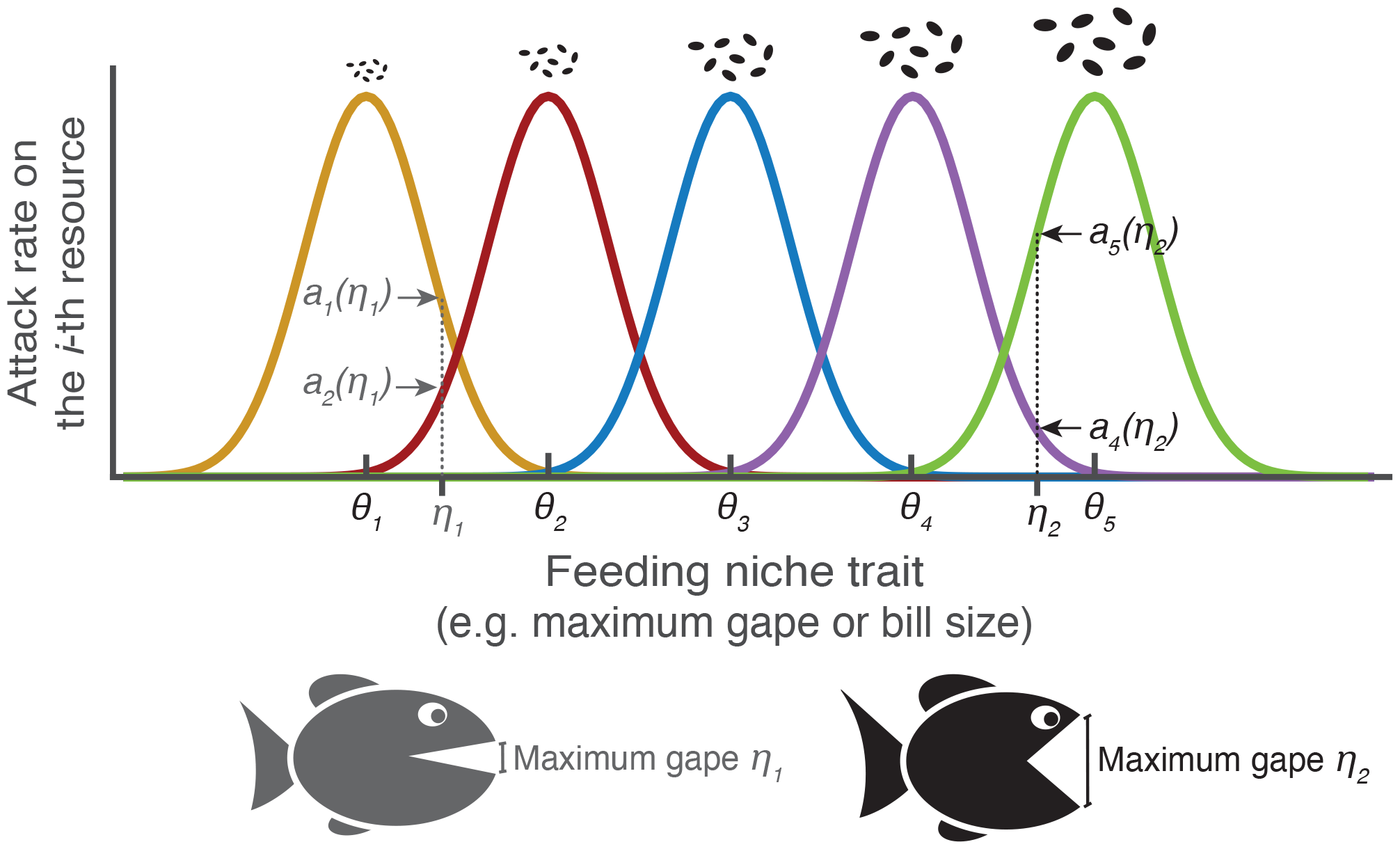
Individual food resource use in the model. Our model considers the existence of multiple niches, or rather the resources that form them, prior to the arrival of an ancestral population. To consume each resource, there exists an optimal feeding niche trait *θ*_*i*_. Organisms differ in an ecological character, i.e. the feeding niche trait (e.g. the maximum gape in fish or reptiles, or the bill size in birds, which determines the size of the food particules that they can ingest). The coloured gaussian curves describe how the attack rate on the *i-*th resource, *a*_*i*_*(η)*, varies with the feeding niche trait *η*. An organism with a small feeding niche trait, e.g. *η*_*1*_ (exemplified by the grey fish) has an attack rate on resource 1 and on resouce 2 indicated by the grey arrows; and its attack rate on other resources is nearly zero. Whereas an organism with a large feeding niche trait, e.g. *η*_*2*_ (exemplified by the black fish) feeds mostly on resource 4 and on resource 5 with attack rates indicated by the black arrows; its attack rate on other resources is nearly zero.

**Figure 2.**
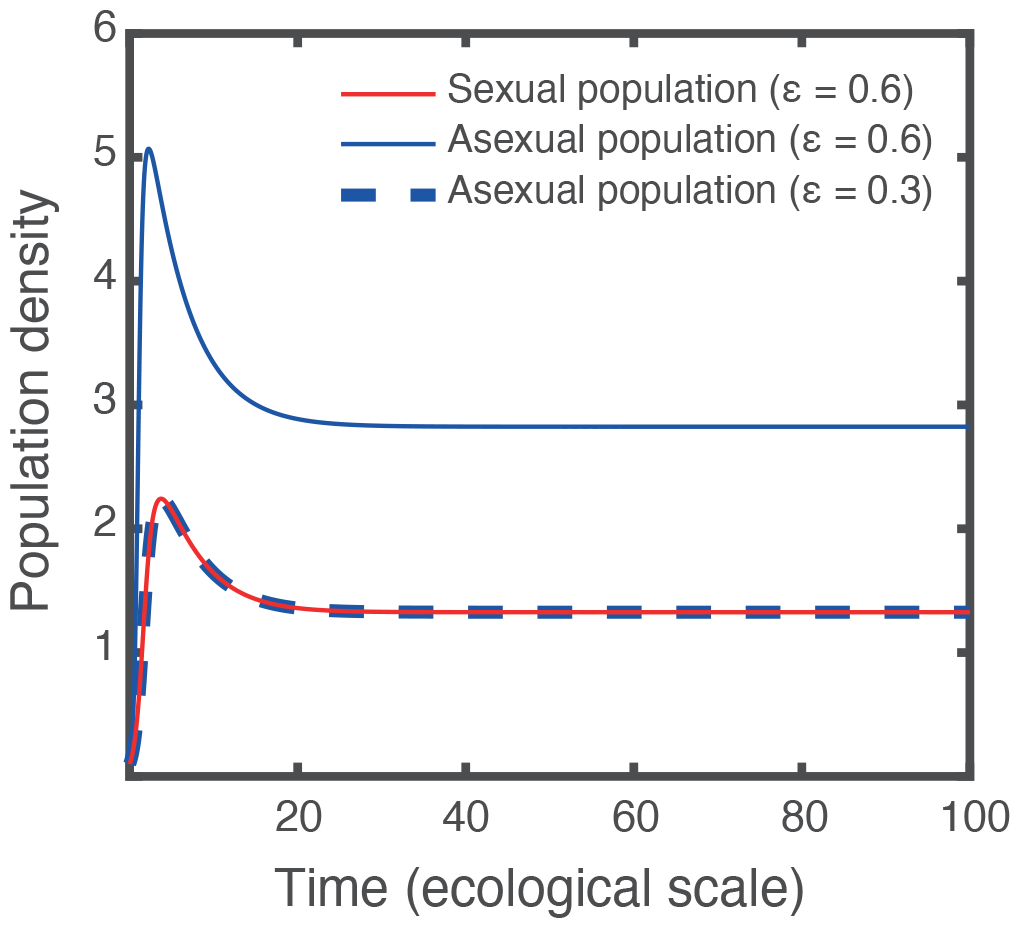
Ecological dynamics of a sexual and asexual population. A sexual population (red line) reaches a lower density than an asexual population (blue solid line) experiencing the same ecological conditions (analytical proof in SI4), and the same density than an asexual population with a half of the efficiency to convert food into offspring (blue dashed line). The ecological dynamics are calculated assuming a (nonevolving) trait value of 1.4, and using eq. 1 and eq. 3 for the sexual population and eq. 2 and eq. 4 for the asexual population. The population encounters two food resources (optimal traits to feed on the resources: *θ*_*1*_= 1 and *θ*_*2*_= 2). The carrying capacity of each food resource is equal to 5 g/L. Other parameter values as in table S1.

#### Individual food resource use and demographic rates

We describe the interaction between consumer individuals and their food resources using a classic Lotka-Volterra predator-prey model. In the model, individuals are characterized by the feeding niche trait *η*, which determines resource use. The attack rate of an individual with trait *η* on the resource *i, a*_*i*_(*η*), equals the maximum attack rate *A* when its feeding niche trait *η* equals *θ*_*i*_, and decreases in a Gaussian manner as *η* moves away from *θ*_*i*_, i.e., *a*_*i*_(*η*) *= A* exp[−(*η* − *θ*_*i*_)^2^/(2*τ*^2^)], where *τ* determines the width of the Gaussian function (figure 1). This implies that there exists a tradeoff to feed on the alternative food resources, such that specialization on one food resource goes at the expense of specialization on the others ^31^. Such tradeoffs have been generally observed in heterotrophic organisms, including bacteria^32^, insects ^33^, and vertebrates ^34,35^. Each individual feeds at a rate *a*_*i*_(*η*)*R*_*i*_ on the *i*th resource, and thus depletes it at this rate (individual food intake thus follows a functional response Type I; see SI1 for functional response Type II). Reproducing individuals convert food into offspring with an efficiency *ε*. Additionally, individuals die at a rate *δ* (mortality is independent of the food intake; see SI1 for food-dependent mortality).

### Eco-evolutionary models

Based on the assumptions described above, we formulate alternative eco-evolutionary models for lineages with sexual and asexual reproduction and use two modelling approaches to analyze them: adaptive dynamics and individual-based models. Adaptive dynamics provides analytical tools to investigate the eco-evolutionary feedback between population dynamics and phenotypic evolution through natural selection ^12,13^. We use these tools to investigate the effect of natural selection alone on diversification. However, adaptive dynamics does not allow for the description of explicit genetic dynamics. Therefore, to investigate the combined effect of natural selection and genetic mixing, we formulate individual-based, genetically explicit models of lineages with sexual and asexual reproduction.

#### Modelling the effect of natural selection alone using adaptive dynamics

We use adaptive dynamics to investigate phenotypic evolution driven only by ecologically-based natural selection. Adaptive dynamics assumes that ecological and evolutionary timescales are sufficiently separated, so that the system reaches the ecological equilibrium before the next phenotypic change occurs.

We first formulate the ecological dynamics of a sexual and an asexual lineage. In the sexual lineage, only females reproduce, producing females and males with equal probability, i.e. primary sex ratio is 1:1. Therefore, we model the ecological dynamics of the sexual lineage considering *l* emerging ecomorphs (*k =* 1, …, *l*) with feeding niche trait *η*_*k*_, female density *F*_*k*_ and male density *M*_*k*_ according to:

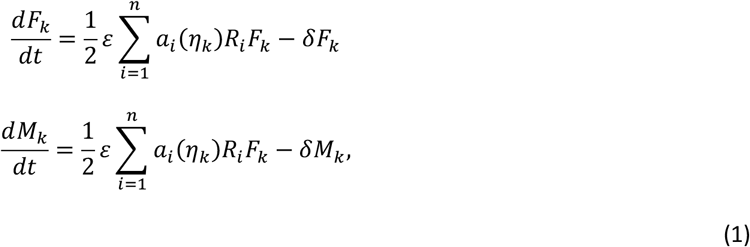

In contrast, in the asexual lineage, all individuals are clonally reproducing individuals. Therefore, the ecological dynamics of the lineage can be modeled considering *m* emerging ecomorphs (*j =* 1, …, *m*) that differ in their feeding niche trait *η*_*j*_. The individual density *N*_*j*_ of each ecomorph follows:

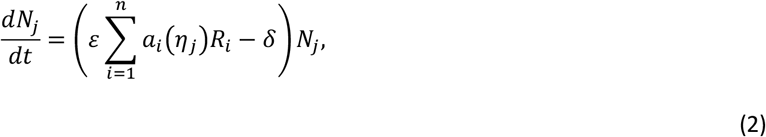

The ecological dynamics of the food resources are given by

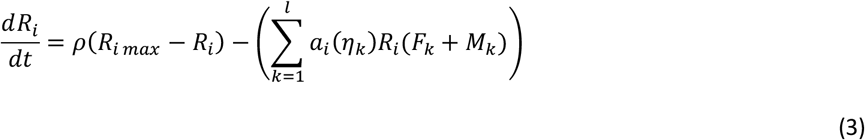

in the case of the sexual lineage, and by

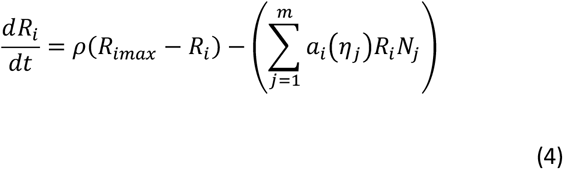

in the case of the asexual lineage.

Based on the ecological dynamics, we then apply adaptive dynamics techniques^36–38^ to study the evolution of the feeding niche trait. Trait change in the sexual lineage is given by:

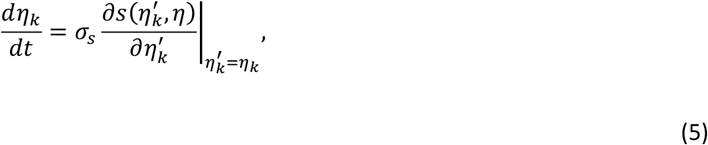

where 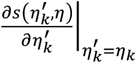 is the selection gradient and *σ*_*s*_ is a constant scaling the evolutionary timescale. Similarly, the trait changes in the asexual lineage at a rate:

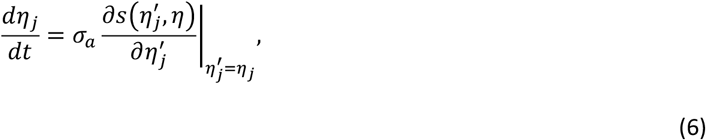

where *σ*_*a*_ is the constant scaling the evolutionary timescale. Diversification occurs through a process of evolutionary branching when directional selection halts, i.e. when 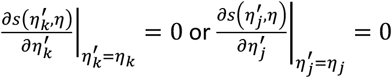, if the trait value at this point corresponds to a minimum of the fitness landscape ^37,38^. Thus, the curvature of the fitness landscape can be used to determine whether a diversification event occurs. The analytical expressions for the selection gradient and the curvature of the fitness landscape are in SI2.

To analyze the effect of natural selection on diversification, we first analytically identify the conditions that enable diversification in an asexual and a sexual population (see SI3). Then, when these conditions are satisfied, we address two questions: 1) do alternative reproductive modes have an effect on the strength of natural selection? and 2) how such effect impacts the rate of evolution in lineages with sexual and asexual reproduction? To address the first question, we analytically compare the selection gradient of two populations differing in their reproductive mode, when each colonizes an environment with two different food resources (see SI5 and figure 3A and 3B). To address the second question, we perform numerical simulations to examine the effect of the reproductive mode on the evolutionary rate of lineages expanding over the trait space to occupy multiple niches (figure 3C). To perform these simulations, the evolutionary dynamics are calculated using eq. 5 and eq. 6, and a constant *σ*_*a*_ = *σ*_*s*_ = 10^−6^ that scales the evolutionary time. Because this constant is used to scale equally the evolutionary time of both sexual and asexual populations, results presented in figure 3C for the lineages with different reproductive mode differ only in the selection gradient, and thus in the strength of natural selection.

**Figure 3.**
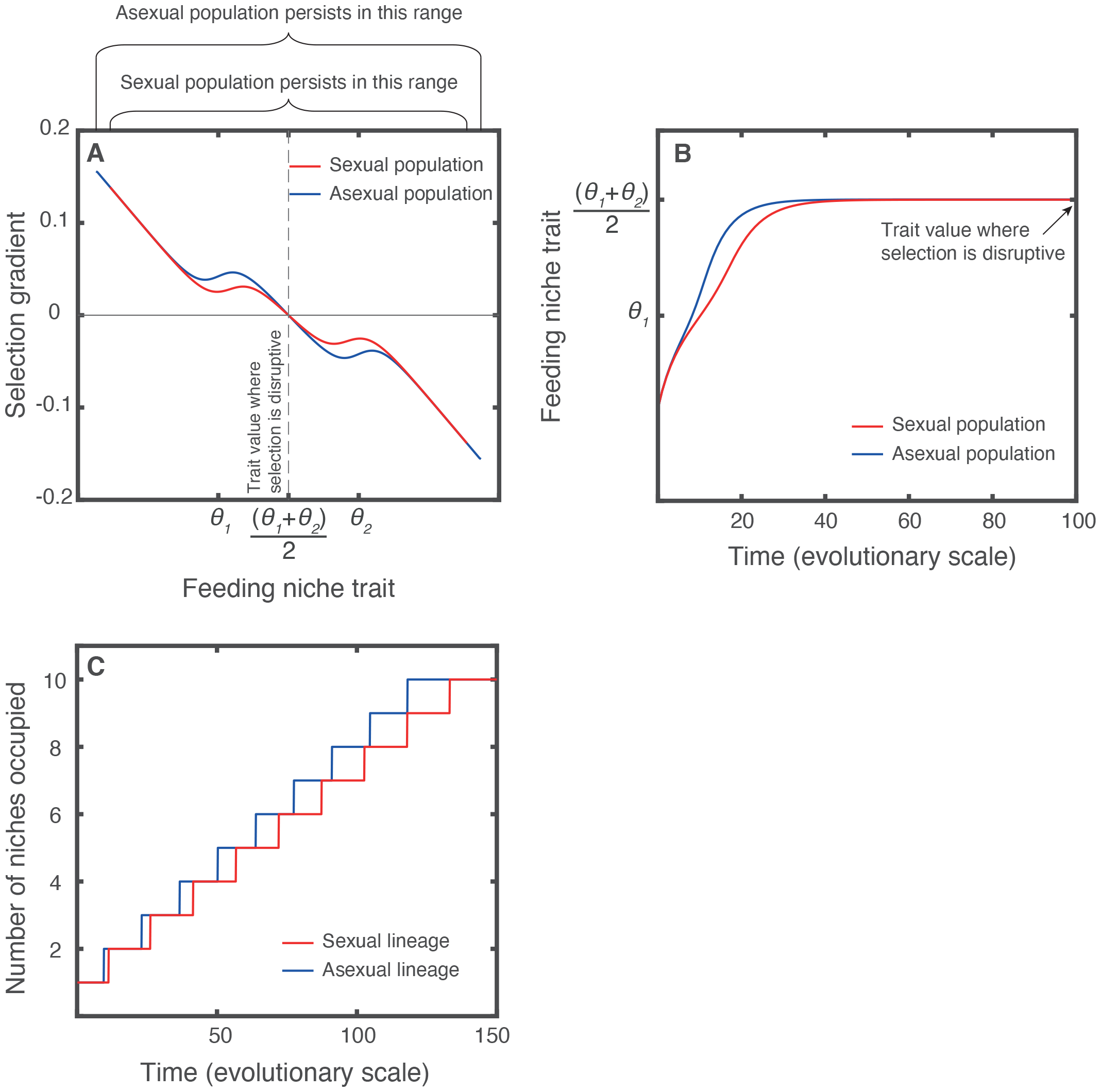
The effect of natural selection alone on diversification. A) Natural selection is stronger in asexual than in sexual populations (proof in SI2), in other words, the magnitude of the selection gradient is always larger in an asexual than in a sexual population experiencing the same ecological conditions. B) Stronger natural selection drives the trait to the value where selection is disruptive faster in an asexual than in a sexual population. C) As a result, in a habitat with multiple niches, an asexual lineage diversifies to occupy all niches faster than a sexual lineage (evolutionary dynamics of both lineages can be found in figure S1). In A and B, a population encounters two food resources (optimal traits to feed on the resources: *θ*_*1*_= 1 and *θ*_*2*_= 2). In C, the ancestral population encounters ten food resources. In all panels, the carrying capacity of each food resource is equal to 5 g/L. Other parameter values as in table S1.

#### Modelling the joint effect of natural selection and genetic mixing using individual-based models

We are interested in understanding the role of genetic mixing in sexual lineages during the diversification process. To do so, we implement genetically explicit individual-based models (IBM) based on the birth, death, and feeding processes as well as the environment described in the section “Model assumptions” (details are in SI6). In the IBM, individual consumers are discrete entities, hence, birth and death occur as discrete, stochastic, events. Because genetic mixing occurs between sexually reproducing individuals, the evolution of barriers to gene flow are required besides ecologically-based disruptive selection for speciation to occur in sexual populations ^1,17^. Following a long tradition in speciation theory, we allow the evolution of such barriers through the evolution of assortative mating ^13,17^. Hereafter, we refer to sexually reproducing individuals as males and females, and to asexually reproducing individuals as clonal individuals. We use an additive diploid multi-locus genetic trait architecture. Each individual is assigned a genotype that determines its phenotype. More precisely, individuals are assigned a set of E-locus that determines its feeding niche trait, such that the sum of all alleles in the E-locus set equals the trait. Alleles of the E-locus set can take any value, and so do the feeding niche trait. Sexually reproducing individuals are assigned an A-locus set that determines the mating trait *ω*, such that the value of the trait equals the average of the allele values, which can be -1 and 1^17^. Assuming that mating depends on female preference only, the alleles in the A-locus set are expressed only in females: females carrying an intermediate mating trait mate randomly (*ω =* 0); females with a negative *ω* mate disassortatively, and females carrying a positive *ω* mate assortatively. Assortability is based on the feeding niche trait and is described by a self-matching mate-choice function^17^. Offspring produce through sexual reproduction inherit parental alleles at each locus independently, in other words, we assume full recombination among all loci. To examine the effect of genetic mixing on the evolutionary rate of a sexual lineage, we vary the contribution of paternal alleles to the offspring’s genes. We do so by assuming that offspring receive in each locus one allele from the father and one from the mother with probability, *p*, or both alleles from the mother with probability, (1 − *p*). When *p =* 1, offspring inherits one maternal and one paternal allele in each locus, meaning that the contribution of each parent is 50% (*p =* 1 is used in all figures, except in figure 5 where it varies). When *p =* 0, offspring inherits only maternal alleles, therefore the contribution of males to offspring is 0%. While biologically unrealistic, the latter scenario represents the case in which natural selection and mutation are the only drivers of evolution. This scenario allows to test whether the results obtained with the adaptive dynamics model are robust to the relaxation of assumptions inherent to this approach, including infinite populations and separation of ecological and evolutionary timescales. In the asexual lineage, we assume clonal apomictic reproduction, therefore offspring inherit the total maternal genetic material (without recombination). Mutation probability is 0.001 per allele (varied in figure S8). When a mutation occurs, the value of the offspring allele is drawn from a normal distribution with a mean equal to the parental value and a standard deviation of *σ*=0.01. To quantify the emergence of novel phenotypes (figure 4D), we divide the feeding niche trait space into bins of width *σ*. A novel phenotype is defined as the first phenotype that emerges within a bin throughout the simulation.

**Figure 4.**
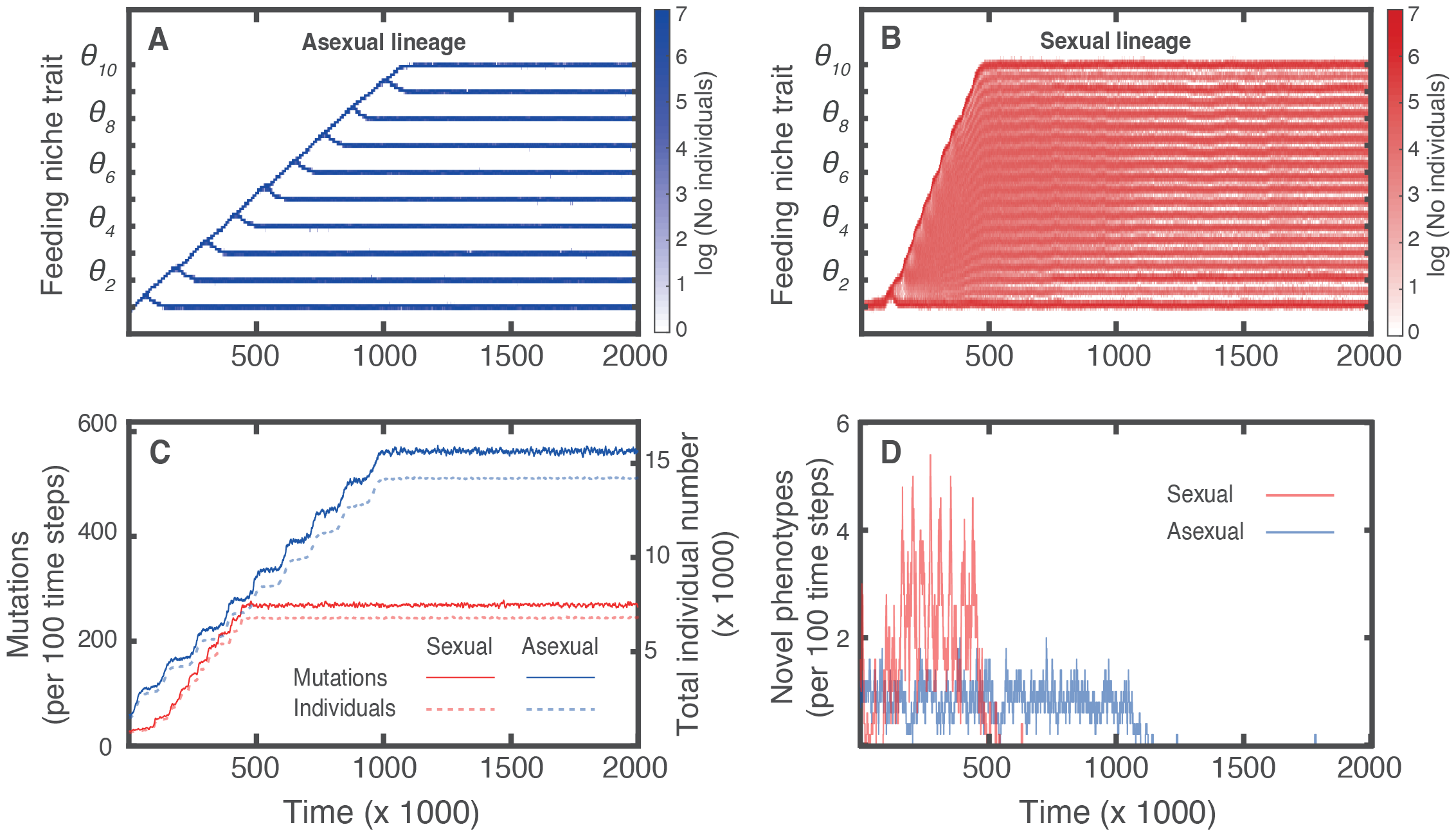
Example simulations of an asexual and a sexual lineage. A) An asexual lineage expands over the trait space through serial diversification events in which one population is split into two. B) A sexual lineage expands over the trait space as a single large population, resembling a hybrid swarm. The diversification process into discrete populations with reduced gene flow occurs later. C) In the asexual lineage, more mutations arise as a consequence of a larger population size. D) During the trait expansion, the rate at which novel phenotypes arise is higher in the sexual lineage. Ticks in the vertical axis in A and B show the trait values corresponding to the optima to feed on food resources. Trait distribution shown in A and B correspond to every 100 time steps in the simulation. Plots in C and D show the moving average of number of individuals, mutations, and novel phenotypes calculated over a sliding window of 50 data points (that is 5000 time steps, since there are 100 time steps between 2 data points). We present the final trait distribution of the lineages in A and B in figure S3. Parameter values as in table S1 and S2.

**Figure 5.**
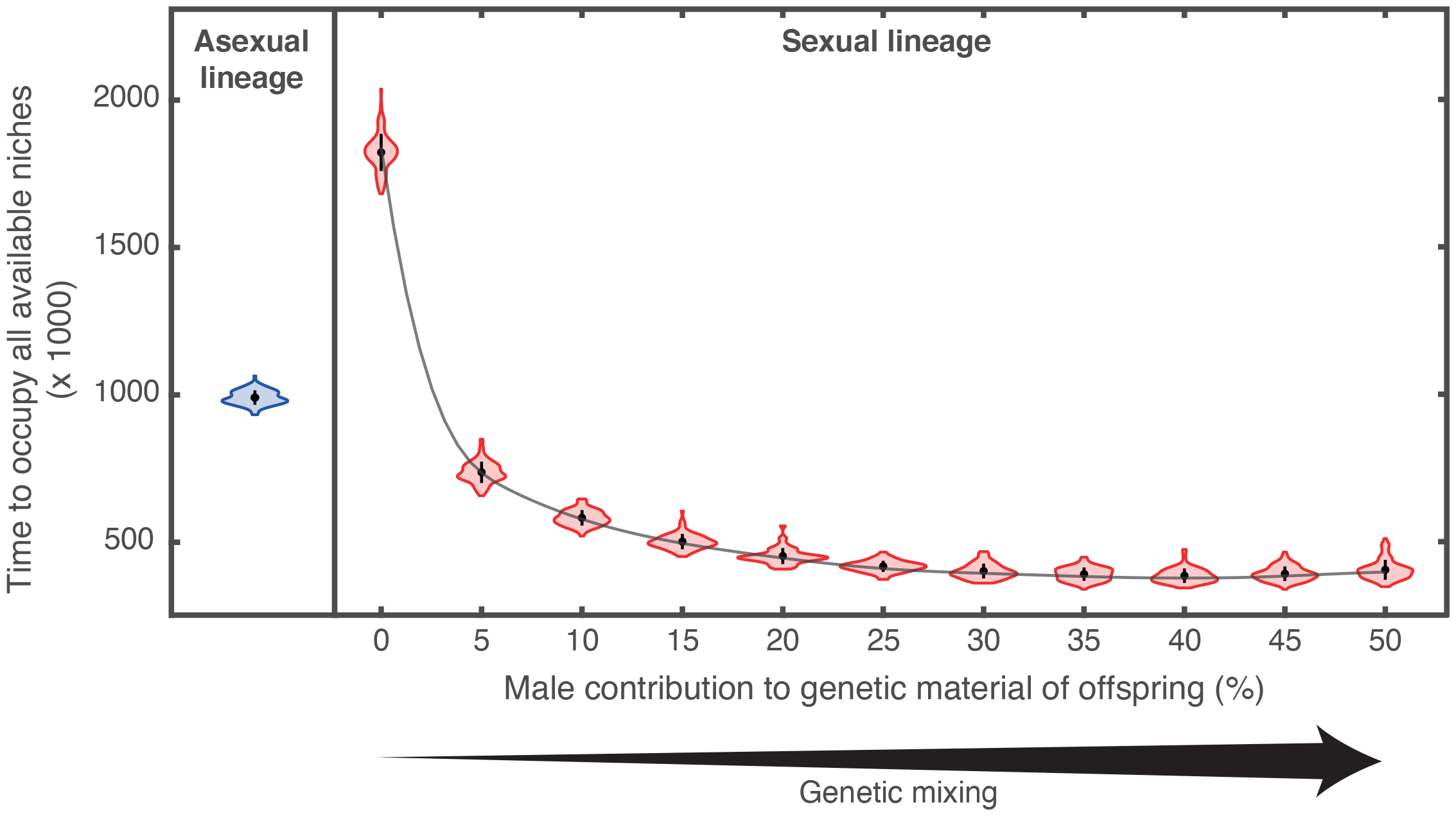
The effect of genetic mixing on the trait expansion process. The expansion over the trait space occurs faster in a sexual lineage with genetic mixing than in an asexual lineage, even when the level of genetic mixing is low. Mean (black dots) and SD (black lines) in each violin plot correspond to 100 replicate simulations. Parameter values as in table S1 and S2.

All models are implemented in MATLAB. Parameterization and initialization of the IBM can be found in SI1 and SI6. Statistical analyses of simulations (figure 5, S4 and S8) were performed using the package ggplot in R. Model implementation code is available in zenodo.

### Robustness analysis

We investigate whether our results are robust to assumptions regarding the functional response Type, a density-independent mortality rate (see SI1 for details), as well as the mutation rate and the number of loci of the feeding niche trait (see SI6 for details). Additionally, we investigate the stability of the attractor of the sexual lineage (final state in figure 4B, see SI7).

## Results

### Conditions that enable diversification in an asexual and a sexual population

Diversification occurs when adaptive evolution can drive a population towards a local minimum of the fitness landscape ^12,13^. At this point, intermediate phenotypes have a fitness disadvantage compared with more extreme phenotypes, and thus selection is disruptive. In a previous work, Chaparro et al. ^28^ derived the conditions for diversification for asexual populations. Here we extend the analysis to the conditions for sexual populations (see analytical derivation in SI3). Two conditions need to be satisfied for diversification to occur in a population colonizing an environment with two different food resources:

- <u>Condition 1</u>: The mean phenotype of the population at an evolutionary equilibrium must be a local fitness minimum. Both an asexual and a sexual population satisfy this condition when (see ref ^28^ and eq. SI3.9 in SI3):

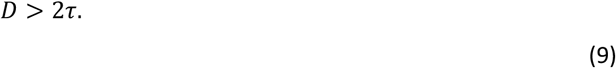

Therefore, when the distance between the optimal trait values to feed on preexisting resources *D* is too small relative to the degree of diet specialization *τ*, disruptive selection does not occur. This is because a sufficiently strong tradeoff between feeding on the alternative food resources is needed to induce disruptive selection ^39,40^. Otherwise, the generalist strategy, corresponding to the evolutionary equilibrium in between the optima to feed on the resources, is a fitness maximum and thus selection is stabilizing.
- <u>Condition 2</u>: This evolutionary equilibrium where the population experiences disruptive selection is an attractor of the evolutionary dynamics, i.e., adaptive evolution can drive a population toward this evolutionary equilibrium. This condition is satisfied when

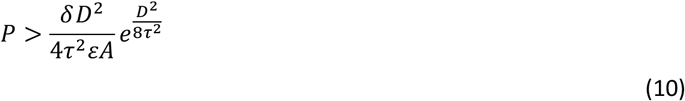

in a population with asexual reproduction ^28^, and when

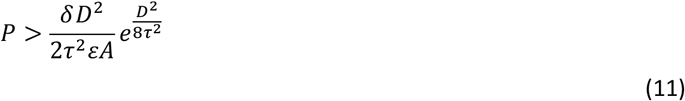

in a population with sexual reproduction (see eq. SI3.25 in SI3). This implies that, for both sexual and asexual populations, there exists a threshold of minimum productivity of the environment *P* that, given other environmental and demographic parameters (*ε, A, δ, τ, D*), enables the trait to be attracted to the point where selection becomes disruptive. This threshold increases with mortality *δ* and decreases with the efficiency to convert food into offspring *ε*. This threshold is higher for a sexual population, meaning that a higher productivity, specifically twice as higher, is required for diversification in a sexual population than in an asexual population experiencing the very same conditions.

Diversification in asexual lineages is less restricted than in sexual lineages, because in asexual populations it can occur at a relatively low level of productivity that is not sufficient to enable diversification in sexual populations. This is the consequence of the reduced reproductive potential of sexual populations. In a sexual population only females, which are only half of the population, have the potential to produce offspring, whereas in an asexual population all individuals can reproduce. As a consequence of the lower reproductive potential, a sexual population has a lower density (figure 2, proof in SI4). Indeed, if the reproductive potential of the asexual population is halved, by for example halving the efficiency to produce offspring, the population density of an asexual population equals that of a sexual population (blue dashed line vs red line in figure 2). As a consequence of this difference in population density, under the very same ecological conditions, competition is stronger among individuals in an asexual than in a sexual population. Because competition is the driver of diversification in our model, stronger competition enables diversification at a lower level of habitat productivity in an asexual population than in a sexual population.

### Natural selection alone drives diversification faster in asexually reproducing lineages

Our study of the selection gradient of an asexual and a sexual population colonizing an environment with two different food resources reveals that natural selection is always stronger in the asexual population (figure 3A, proof in SI5). This is because, under the same ecological conditions, an asexual population has a higher density than a sexual population. Higher density leads to stronger competition among individuals in an asexual population, causing stronger natural selection. As a result, if the evolutionary rate is driven only by natural selection, evolution drives the feeding trait of an asexual population to the value where selection is disruptive faster than it drives the trait of a sexual population (figure 3B). In scenarios with several niches, natural selection therefore causes an asexual lineage to diversify and occupy the available niches faster than a sexual lineage (figure 3C). A further analysis reveals that competition is stronger in an asexual than in a sexual population when considering, not only the birth, but also the mortality rate to be dependent on the feeding rate (figure S2). In conclusion, when natural selection is the only driver of evolution, the diversification process is faster in an asexual lineage than in a sexual lineage.

### Natural selection and genetic mixing jointly drive diversification faster in sexually reproducing lineages

We next consider the combined effect of genetic mixing and natural selection. Individual-based simulations of lineages with different reproductive mode indicate that sexual lineages fill up niches faster than asexual lineages (figure 4), opposite to what we observed when natural selection acts alone (figure 3C). The expansion over the trait space in an asexual lineage occurs through sequential diversification. The ancestral population evolves towards the trait value in between the optimum to feed on resource 1 and the optimum to feed on resource 2 (in between *θ*_1_ and *θ*_2_ in figure 3A). At this point, the population experiences disruptive selection and undergoes a diversification event. After diversification, the traits of the two resulting populations diverge. One of the populations evolves towards the trait value that is optimal to feed on resource 1, whereas the other evolves towards the trait value in between the optima to feed on resources 2 and 3. The first population now experiences stabilizing selection. The second population, in contrast, experiences disruptive selection again, resulting in a new diversification event. This alternation between adaptation through directional selection and diversification through disruptive selection is repeated until all preexisting resources are fully utilized by the lineage. As a result of this process, the asexual lineage diversifies into the same number of populations as the number of existing niches.

Conversely, the expansion over the trait space in a sexual lineage occurs without diversification. A single sexual population rapidly expand over the trait space, filling up all available niches in less than half of the time that the asexual lineage takes to do so (figure 4B; this faster expansion of a sexual lineage occurs independently of the functional response type for individual food intake, see figure S4). Assortative mating evolves early during the trait expansion (figure S5), while diversification occurs much later. Discrete clusters emerge gradually, some are specialists (their mean trait is approximately equal to some optimum) and others are generalists (their mean trait is approximately equal to the intermediate value between two optima). By the end of the simulation, these clusters are differentiated and individuals mate assortatively based on their feeding niche trait, resulting in gene flow below 5% or a level of reproductive isolation above 0.9 (based on statistics^41^). At the end of the simulation, the sexual lineage has a larger diversity, partly due to the coexistence of specialists and generalists. Further analyses suggest that this state in which specialists and generalists coexist is stable (see SI7). Despite the formation of discrete populations occurs much later in the sexual lineage, the faster expansion over the trait space enables an earlier occupancy of available niches for this lineage. The faster trait expansion of sexual lineages is mostly explained by the higher rate at which novel phenotypes are generated (figure 4D), which occurs despite less mutations take place due to the smaller population size (figure 4C). This process does not result immediately in a more specious lineage because diversification occurs much later in the sexual lineage. However, in the long run, an environment colonized by a sexual and an asexual lineage with the same history will be dominated by sexual species due to the capacity of the sexual lineage to fill up niches faster (figure S6A), even with a small amount of genetic mixing. In the face of disturbances, the faster niche occupancy enables a sexual lineage to rapidly replace extinct populations by novel populations emerging through diversification (figure S6B). As a result, sexual lineages can readily recover their pre-disturbance diversity, or even increase it by further expanding over the trait space, occupying vacant niches left by extinct asexual populations (see SI7).

### A small amount of genetic mixing is sufficient to overturn the effect of natural selection

To further assess the contribution of genetic mixing to the trait expansion and diversification process, we ran individual-based simulations of the sexual lineage varying the level of genetic mixing. Without genetic mixing, a sexual lineage takes very long time to fill up available niches, almost double of the time that an asexual lineage takes to do so (figure 5). Because in the absence of genetic mixing the main force driving evolution is natural selection, this result supports our finding, from adaptive dynamics models, that natural selection alone promotes faster diversification in asexually reproducing lineages. Conversely, with a small amount of genetic mixing, the time required for the sexual lineage to occupy all available niches is significantly reduced. This reduction in time required to fill in all available niches as a consequence of increasing genetic mixing behaves in a nonlinear manner: small increases in male contribution to offspring’s genetic material have a larger impact when genetic mixing is low (red violins at the left in figure 5) than when it is high (violins at the right). Even relatively low genetic mixing can speed up the trait expansion process in a sexual lineage to enable a faster niche occupancy than that of an asexual lineage.

## Discussion

Our results reveal that the reproductive mode substantially affects ecological diversification. We show that sexual reproduction leads to faster diversification. Similarly, Melian et al. ^23^ found that sexual reproduction increased speciation rates in a neutral model. However, in this study, lineages with sexual reproduction had lower species richness because extinction rates exceeded speciation rates. Our simulations show that sexual populations are smaller than asexual populations (figure 4C), which increases their extinction risk. However, the higher risk of extinction can be balanced by a faster speciation process (figure S6 and S7B). Therefore, our findings provide support for the emergence of more specious lineages associated to sexual reproduction. In line with these findings, a recent phylogenetic analysis showed that lineages with sexual reproduction have accelerated rates of diversification (relative to asexual lineages), and that patterns of species richness are strongly related to these differences in diversification rates ^7^. This statistically based analysis therefore supports our findings that sexual reproduction is associated with faster diversification and higher species richness.

Going a step further, we investigate how natural selection and genetic mixing influence ecological diversification in lineages with sexual and asexual reproduction. We find diversification to be less restrictive in asexual populations (SI3) and natural selection to be stronger during the diversification process in asexual lineages (figure 3, proof in SI5). Both findings are the consequence of the paradox of sex described by Maynard-Smith ^18,22^; he acknowledged that sexual reproduction has an immediate cost relative to asexual reproduction because sexual females invest half their reproductive resources into males that, in turn, invest minimally into each offspring sired. Because asexuality is exactly twice as efficient at converting resources into offspring, intraspecific competition is stronger in an asexual population. Competition drives natural selection, therefore the twofold cost of sex translates into weaker selective pressures in sexual populations that, on one hand, can hinder diversification, for example when resource productivity is low, and on the other hand, can slow down the diversification process. We provide evidence that genetic mixing can overturn the effect of selection alone, speeding up the emergence of novel phenotypes and thus the trait expansion process in sexual lineages (figure 4). Despite weaker selective pressures, trait expansion is therefore faster in sexual than in asexual lineages, and this result is robust to variation in mutation rate and the number of loci encoding the feeding niche trait (figure S8). This effect is complementary to the observations that sex speeds adaptation by allowing natural selection to more efficiently bring together mutations that confer a fitness advantage ^42–46^.

Our results show a strong nonlinear effect of genetic mixing. A small amount of genetic mixing in sexual organisms is needed to accelerate the trait expansion of a lineage beyond the speed of an analogous lineage of asexual organisms (figure 5). Furthermore, increases in genetic mixing when its level is already high do not have a significant effect on the speed of the trait expansion. A similar nonlinear effect of genetic mixing has been observed when organisms occasionally engage in sex, such as in species with facultative sex. In these species, the evolutionary rate can be nearly as fast when the frequency of sex is low than when it occurs in every reproductive event ^47–49^. By reducing the frequency of sex and thus the rate of male production, these species receive the benefits of sex while paying a lower cost than the twofold cost of sex. In our model, sex is obligate and females produce daughters and sons in equal proportion. An extension of our model implementing facultative sex might increase the efficiency at converting resources into offspring (relative to a population with obligate sex). This could strengthen natural selection and thus speed up diversification. Hence, a lineage with facultative sex could have a faster diversification rate than an obligate sexual lineage. Theory has suggested that facultative reproductive strategies should be more successful than obligate sexual or asexual strategies for adaptation (microevolution) ^47–49^. Our findings suggest that this may be the case as well for diversification (macroevolution).

In the asexual lineage, we model reproduction through apomixis, i.e., offspring inherit the total maternal genetic material without recombination. This reproductive system is common in many bacteria and fungi but also in some animals such as aphids and snails ^50,51^. There are also organisms that reproduce asexually via automixis, in which a modified meiosis takes place in asexual females. In this case, offspring develop from unfertilized but diploid eggs and may be genetically diverse and distinct from the mother. Such reproductive system is present in several animal taxa including arthropods, nematods, moluscs and vertebrates^52^. Despite the lack of genetic mixing among individuals, automictic populations are expected to have a higher genetic variability than apomictic populations ^52^. We would therefore expect that a lineage with automictic reproduction will diversify faster than a lineage with apomictic reproduction, but probably slower than a lineage with sexual reproduction. Further research will be required to know how much can automixis speed up diversification in an asexual lineage.

Similar to classic models of ecological diversification ^17,53^, in our model, gene flow between phenotypically divergent clusters of sexual organisms is reduced by the evolution of assortative mating based on the ecological character, in our case the feeding niche trait. There is growing evidence of the existence of characters that influence both mating patterns and ecological fitness in a variety of taxa, including plants, vertebrates, and invertebrates^54^. Therefore, assortative mating mechanisms such as the one modeled here may be common in nature, in particular when their associated costs are low. Conversely, when assortative mating entails high costs, genetic variation for mate choosiness can be significantly reduced, preventing speciation ^24,55^.

Our findings may be relevant to understand the paradox of sex, often stated as: why a larger number of eukaryotic species reproduce sexually relative to those that reproduce asexually despite the costs of sex? Most research addressing this problem has focused on identifying multiple mechanisms that enable the maintenance of sex within populations^19,48,56–74^. However, differences in diversification rate may also contribute to the relative richness of species with each reproductive mode. Our results suggest that sexual reproduction may also be widespread among species because it increased rates of diversification. Our results place the well-known long-term benefits of sex ^42–46^ in a macroevolutionary context that can help explain broad diversity patterns associated to each reproductive mode. Yet, future work is required to link them to the short-term benefits of sex that enable the sexual individuals to avoid being outcompeted by asexual ones. The integration of the well-known mechanisms for the maintenance of sex within populations with ecological diversification might further explain the widespread occurrence of sexual reproduction. We show how such integration can be done using models that link macroevolutionary patterns to microevolutionary and ecological processes.

## Acknowledgements

We thank Jukka Jukela, Hanna ten Brink and Hanna Kokko for valuable discussions and helpful comments on an earlier version of this paper. This research was funded by the European Union’s Horizon 2020 research and innovation programme under the Marie Skłodowska-Curie grant agreement no. 898687 awarded to CCP.

## Author contributions

CCP conceived the idea. CCP and GR analyzed the model. CCP led the writing of the manuscript, and CM and GR critically revised and contributed to later versions. All authors gave final approval of the manuscript to be published.

