## Supplementary material for "Ecological diversification in sexual and asexual lineages"

**Supplementary information for:**

### Genetic mixing, diversification and the widespread occurrence of sex in nature

This supplementary note includes:

- Supplementary text:
    1. Model parameterization and robustness analyses
    2. Analytical derivation of the fitness gradient and curvature of the fitness landscape
    3. Analytical derivation of conditions that enable diversification in a sexual population
    4. Analytical proof that the density of an asexual population is higher than the density of a sexual population at ecological equilibrium
    5. Analytical investigation of the strength of selection in populations with different reproductive modes
    6. Individual-based model details
    7. Investigation of stability of the final state of a sexual lineage
- Figures S1 to S10
- Table S1 in SI1 and S2 in SI6

Other supplementary materials for this manuscript includes: MATLAB code of the model for numerical simulations and the individual-based models.

### Supplementary Information 1. Model parameterization and robustness analyses

***Model parameterization***

In a previous study, Chaparro et al.(1) showed that the selection of the values for the parameters $\varepsilon, A, \delta, D, \tau, P$ is crucial to enable diversification. Following this work, we assign the values used in this study to the demographic parameters $\varepsilon, A, \delta$. Then, we select the values of the degree of specialization, $\tau$, and the productivity, $P$, such that the conditions for diversification in sexual populations are satisfied given that the distance between optima to feed on preexisting resources $D = 1$ (analytical derivation can be found SI3).

Parameter values are summarized in table S1.

Table S1. Variables and parameters

| Variable or parameter | Symbol | Value | Units |
|---|---|---|---|
| Variables | | | |
| Feeding niche trait | $\eta$ | Evolving trait | - |
| Individual density | $N_j$ | - | $L^{-1}$ |
| Female density** | $F_k$ | - | $L^{-1}$ |
| Male density** | $M_k$ | - | $L^{-1}$ |
| Food resource density | $R_i$ | - | g $L^{-1}$ |
| Environmental parameters | | | |
| Renewal rate of resources† | $\rho$ | 0.01 | (unit of time)$^{-1}$ |
| Total productivity of the habitat | $P$ | 50 | g $L^{-1}$ |
| Number of resources | $n$ | 10 | - |
| Distance between optima to feed on the resources | $D$ | 1 | - |
| Demographic parameters | | | |
| Maximum attack rate† | $A$ | 0.2 | L (unit of time)$^{-1}$ |
| Width of the Gaussian curve describing the degree of specialization to successfully attack a food resource | $\tau$ | 1/3 | - |
| Assimilation efficiency | $\varepsilon$ | 0.6 | - |
| Mortality rate† | $\delta$ | 0.02 | (unit of time)$^{-1}$ |

†Rates have an arbitrary unit of time
**Variables or parameters apply only to sexual population(s)

***Robustness analysis***

We furthermore investigate whether our results are robust to different assumptions regarding the functional response Type (figure S4). We do so by using a Type II functional response, such that the food intake of an individual with trait $\eta_j$ is:

$$\frac{\sum_{i=1}^{n} a_i(\eta_j) R_i}{1 + h_T\left(\sum_{i=1}^{n} a_i(\eta_j) R_i\right)},$$

(SI1.1)

where, $h_T$ is the handling time.

We additionally investigate whether our results are robust to relaxing the assumption that individuals die at a rate that is independent of the food consumption (figure S2). To do so, we define a maximum death rate, $\delta_{\max}$, that individuals experience when their feeding rate, $\varphi$ ($\varphi = \sum_{i=1}^{n} a_i(\eta_k) R_i$ in a sexually reproducing lineage and $\varphi = \sum_{i=1}^{n} a_i(\eta_j) R_i$ in an asexually reproducing lineage), approaches zero, and a minimum death rate, $\delta_{\min}$, that individuals experience when their feeding rate equals their maximum possible feeding rate, or in other words, their feeding rate under conditions of replete food densities, i.e. $\varphi|_{R_i=R_{i\,\max}}$. For the sake of simplicity, we assume that the mortality rate changes linearly between the maximum and the minimum (figure S2A). Hence, the ecological dynamics follow

$$\frac{dF_k}{dt} = \frac{1}{2}\varepsilon \sum_{i=1}^{n} a_i(\eta_k) R_i F_k - \delta_d F_k$$

$$\frac{dM_k}{dt} = \frac{1}{2}\varepsilon \sum_{i=1}^{n} a_i(\eta_k) R_i F_k - \delta_d M_k,$$

(SI1.2)

for a sexually reproducing lineage, and

$$\frac{dN_j}{dt} = \left(\varepsilon \sum_{i=1}^{n} a_i(\eta_j) R_i - \delta_d\right) N_j,$$

(SI1.3)

for an asexually reproducing lineage. In eq. SI1.2 and SI1.3, $\delta_d = \delta_{\max} + \varphi \frac{\delta_{\min} - \delta_{\max}}{\varphi|_{R_i=R_{i\,\max}}}$.

### Supplementary Information 2. Analytical derivation of the selection gradient and curvature of the fitness landscape

Here, we derive the selection gradient and the curvature of the fitness landscape using the adaptive dynamics framework of an asexual and a sexual lineage. According to adaptive dynamics theory, these quantities depend on the invasion fitness, which is the relative fitness of a rare mutant population in the environment set by (resident) ecomorph populations with trait values $\eta = (\eta_1, \dots, \eta_m)$.

**Sexual lineage** To model the evolutionary dynamics of the sexual lineage, we follow the adaptive dynamics method developed by Metz and de Kovel for diploid organisms with dioicous sexual reproduction. According to this method, the lifetime reproductive output, denoted here by $L$, can be used as a proxy for fitness. Therefore, the invasion fitness of a rare mutant with feeding niche trait $\eta_{k'}$ in the environment set by resident populations with trait values $\eta = (\eta_1, \dots, \eta_l)$ equals

$$L(\eta_{k'}, \eta) = \frac{1}{2}\left(f_F \ell_F(\eta_{k'}, \eta) + f_M \ell_M(\eta_{k'}, \eta)\right) \tag{SI2.1}$$

where $f_F$and $f_M$ are the fractions of newly produced females and males, and $\ell_F$ and $\ell_M$ are the average lifetime numbers of offspring begotten by a female or male, respectively. Because in our model females and males are produced in equal proportion, then $f_F = f_M = 1/2$. Similarly, the average lifetime numbers of offspring begotten by females and males are equal because they do not differ in any of their demographic traits. Thus $\ell_F(\eta_{k'}, \eta) = \ell_M(\eta_{k'}, \eta) = \varepsilon/\delta \sum_{i=1}^{n} a_i(\eta_{k'}) R_i^*(\eta)$, where $R_i^*(\eta)$ corresponds to the resource food densities at ecological equilibrium. According to the adaptive dynamics approach, in the sexual lineage the rate of change of the feeding niche trait of the $k$th ecomorph population can be approximated by:

$$\frac{d\eta_k}{dt} = \sigma_s \frac{\partial L(\eta_{k'}, \eta)}{\partial \eta_{k'}} \tag{SI2.2}$$

Because we use $L$ as proxy for fitness, the fitness gradient $\frac{\partial s(\eta_{k'}, \eta)}{\partial \eta_{k'}}$ found in eq. 5 in Methods is replaced by $\frac{\partial L(\eta_{k'}, \eta)}{\partial \eta_{k'}}$.

From eq. SI2.1, we obtain

$$\frac{\partial L(\eta_{k'}, \eta)}{\partial \eta_{k'}} = \frac{\varepsilon}{2\delta} \sum_{i=1}^{n} A \frac{(\theta_i - \eta_{k'})}{\tau^2} e^{\frac{-(\theta_i - \eta_{k'})^2}{2\tau^2}} R_i^*(\eta) \tag{SI2.3}$$

Directional selection halts where the selection gradient vanishes, in other words when equation (SI2.3) equals 0. At this point, selection can be stabilizing if this trait value corresponds to a fitness maximum, or disruptive if it corresponds to a fitness minimum. The curvature of the fitness function therefore determines whether evolution halts or an ecomorph splits into two different ecomorphs. Hence, the second derivative of the fitness function with respect to $\eta_{k'}$ evaluated at the ecological equilibrium when resident ecomorphs have traits $\eta$ allows us to determine whether a diversification event occurs:

### Supplementary Information 3. Analytical derivation of conditions that enable diversification in a sexual population

Here, we derive the conditions that enable diversification when one sexual population with female density $F$ and male density $M$ colonizes an environment with two different food resources with densities $R_1$ and $R_2$. For diversification to occur, a local minimum of the fitness landscape must be an attractor of the evolutionary dynamics. Thus, two conditions must be fulfilled for diversification to occur: 1) the mean phenotype of the population at evolutionary equilibrium corresponds to a local fitness minimum such that selection is disruptive, and 2) this equilibrium is an attractor of the evolutionary dynamics. We restrict our analytical investigation of these conditions to the simplest case, in which $R_{1max} = R_{2max} = R_{max}$. The optimal niche trait values to feed on resource 1 and resource 2 are $\theta_1$ and $\theta_2$, respectively, and their distance in the trait axis equals $D$. Without a loss of generality, we assume that $\theta_1 = 0$ and $\theta_2 = D$. Following the adaptive dynamics framework, we assume that evolution occurs much more slowly than the ecological dynamics. This separation of the ecological and evolutionary timescales enables us to approximate the ecological dynamics by assuming that they are always in a quasi-steady state determined by the ecomorph population with trait value $\eta$. We can thus derive analytical expressions for the population and resource densities at ecological equilibrium (see derivation of these expressions in Ecological equilibrium of an asexual and a sexual population in SI4). Subsequently, using these expressions, we investigate the fitness landscape of a rare mutant in the environment set by the population with a trait value $\eta$.

**1. When does a population experience disruptive selection?** One of the necessary conditions for diversification is that a population experiences disruptive selection. This occurs when the population is in an evolutionary equilibrium that corresponds to a fitness minimum, hence the curvature of the fitness landscape around this trait value is positive. To determine when this condition is satisfied, we investigate the fitness landscape of a rare mutant with niche trait value, $\zeta$, in the environment set by the population with a trait value, $\eta$. The fitness experienced by a mutant is given by

$$W(\zeta, \eta) = \frac{1}{2}\varepsilon\left(a_1(\zeta)R_1(\eta) + a_2(\zeta)R_2(\eta)\right) - \delta. \tag{SI3.1}$$

and thus its fitness gradient is

$$\begin{aligned} \frac{\partial W}{\partial \zeta}(\eta, \eta) &= \frac{1}{2}\varepsilon\left(\frac{\partial a_1}{\partial \zeta}R_1(\eta) + \frac{\partial a_2}{\partial \zeta}R_2(\eta)\right) \\ &= -\frac{\varepsilon}{2\tau^2}\eta(a_1(\eta)R_1(\eta) + a_2(\eta)R_2(\eta)) + \frac{\varepsilon D}{2\tau^2}a_2(\eta)R_2(\eta) \\ &= -\frac{\delta}{\tau^2}\eta + \frac{\varepsilon D}{2\tau^2}a_2(\eta)R_2(\eta). \end{aligned} \tag{SI3.2}$$

A population with a trait value $\eta^*$ is in an evolutionary equilibrium if, at this point, its selection gradient equals zero, i.e. $\frac{\partial W}{\partial \zeta}(\eta^*, \eta^*) = 0$. This equilibrium corresponds to a fitness minimum if $\frac{\partial^2 W}{\partial \zeta^2}(\eta^*, \eta^*) \geq 0$. To evaluate these conditions when $\eta^* = \frac{D}{2}$, we first need to calculate $\frac{\partial^2 W}{\partial \zeta^2}(\eta, \eta)$. Derivating equation (SI3.2), we have

$$\begin{aligned}
\frac{\partial^2 W}{\partial \zeta^2}(\eta,\eta) &= \frac{1}{2}\varepsilon\left(\frac{\partial^2 a_1}{\partial \zeta^2}R_1(\eta) + \frac{\partial^2 a_2}{\partial \zeta^2}R_2(\eta)\right)\\
&= \frac{1}{2}\varepsilon\left(\frac{1}{\tau^2}a_1(\eta)\left(\frac{\eta^2}{\tau^2}-1\right)R_1(\eta) + \frac{1}{\tau^2}a_2(\eta)\left(\frac{(\eta-D)^2}{\tau^2}-1\right)R_2(\eta)\right)\\
&= \frac{\varepsilon}{2\tau^2}\left((\frac{\eta^2}{\tau^2}-1)(a_1(\eta)R_1(\eta)+a_2(\eta)R_2(\eta)) + \frac{D}{\tau^2}a_2(\eta)R_2(\eta)(D-2\eta)\right)
\end{aligned} \tag{SI3.3}$$

When $\eta^* = \frac{D}{2}$, $R_1(\frac{D}{2}) = R_2(\frac{D}{2})$, hence

$$\varepsilon a_2(\frac{D}{2})R_2(\frac{D}{2}) = \delta \tag{SI3.4}$$

because in any ecological equilibrium the per capita growth rate equals 0 (i.e. $W(\eta, \eta) = 0$). Using equations (SI3.2) and (SI3.4), we can show that $\eta^* = \frac{D}{2}$ is an evolutionary equilibrium for any $D > 0$.

$$\frac{\partial W}{\partial \eta}(\frac{D}{2},\frac{D}{2}) = -\frac{\delta}{\tau^2}\frac{D}{2} + \frac{\varepsilon D}{2\tau^2}a_2(\frac{D}{2})R_2(\frac{D}{2}) \tag{SI3.5}$$

$$= \frac{D}{2\tau^2}(-\delta + \varepsilon a_2(\frac{D}{2})R_2(\frac{D}{2})) \tag{SI3.6}$$

$$= 0 \tag{SI3.7}$$

Finally, we evaluate the curvature of the fitness landscape at $\eta^* = \frac{D}{2}$ using equations (SI3.3) and (SI3.4)

$$\frac{\partial^2 W}{\partial \zeta^2}(\frac{D}{2},\frac{D}{2}) = \frac{\delta}{\tau^2}(\frac{D^2}{4\tau^2}-1) \tag{SI3.8}$$

The curvature of the fitness landscape for $\eta^* = \frac{D}{2}$ is positive, i.e. $\frac{\partial^2 W}{\partial \zeta^2}(\frac{D}{2}, \frac{D}{2}) \geq 0$, and thus selection is disruptive in the evolutionary equilibrium $\eta^* = \frac{D}{2}$ when

$$D > 2\tau \tag{SI3.9}$$

**2. When does a population evolve toward the evolutionary equilibrium where selection is disruptive?** The second necessary condition for diversification is that a population evolves toward the evolutionary equilibrium where selection is disruptive. This occurs when this equilibrium is an evolutionary attractor. To determine for which values of $D$, the equilibrium $\eta^* = \frac{D}{2}$ is an attractor, we investigate its stability.

An evolutionary equilibrium $\eta^*$ is stable if $\frac{\partial}{\partial \eta}\left(\frac{\partial W}{\partial \zeta}(\eta,\eta)\right) < 0$. Hence, we first derivate equation (SI3.2) with respect to $\eta$.

$$\frac{\partial}{\partial \eta}\left(\frac{\partial W}{\partial \zeta}(\eta,\eta)\right) = -\frac{\delta}{\tau^2} + \frac{\varepsilon D}{2\tau^2}\frac{\partial a_2}{\partial \eta}(\eta)R_2(\eta) + \frac{\varepsilon D}{2\tau^2}a_2(\eta)\frac{\partial R_2}{\partial \eta}(\eta) \tag{SI3.10}$$

Next, we calculate $\frac{\partial a_2}{\partial \eta}(\eta)$

$$\frac{\partial a_2}{\partial \eta}(\eta) = a_2(\eta)\frac{D-\eta}{\tau^2}, \tag{SI3.11}$$

and $\frac{\partial R_2}{\partial \eta}(\eta)$ by derivating eq. (SI4.19) with respect to $\eta$

$$\begin{aligned}\frac{\partial R_2}{\partial \eta} &= \frac{-\rho R_{max}(\frac{\partial a_2}{\partial \eta}2F + a_2\frac{\partial F}{\partial \eta})}{(2a_2F+\rho)^2} \\ &= -R_2\frac{\frac{\partial a_2}{\partial \eta}2F + a_2\frac{\partial F}{\partial \eta}}{2a_2F+\rho}\end{aligned}$$

$\frac{\partial F}{\partial \eta}$ can be obtained by derivating eq. (SI4.21), (SI4.22), (SI4.23) and (SI4.24):

$$\frac{\partial F^*}{\partial \eta} = \frac{2\alpha_s\left(-\frac{\partial \beta_s}{\partial \eta} + \frac{\partial}{\partial \eta}(\sqrt{\Delta_s})\right) - \left(-\beta_s + \sqrt{\Delta_s}\right)\frac{\partial \alpha_s}{\partial \eta}}{4\alpha_s^2} \tag{SI3.12}$$

where

$$\frac{\partial \alpha_s}{\partial \eta} = \frac{\delta}{\tau^2}a_1(\eta)a_2(\eta)(D-2\eta) \tag{SI3.13}$$

$$\frac{\partial \beta_s}{\partial \eta} = -\frac{\rho\delta\eta}{\tau^2}a_1(\eta) - \frac{\rho\delta(\eta-D)}{\tau^2}a_2(\eta) - \frac{\varepsilon\rho R_{max}}{\tau^2}a_1(\eta)a_2(\eta)(D-2\eta) \tag{SI3.14}$$

$$\frac{\partial\sqrt{\Delta_s}}{\partial \eta} = \frac{1}{2\sqrt{\Delta}}\left(2\delta^2\rho^2(a_1(\eta)-a_2(\eta))(\frac{\partial a_1}{\partial \eta} - \frac{\partial a_2}{\partial \eta}) + 2\rho^2\varepsilon^2R_{max}^2a_1(\eta)a_2(\eta)(a_2(\eta)\frac{\partial a_1}{\partial \eta} + a_1(\eta)\frac{\partial a_2}{\partial \eta})\right) \tag{SI3.15}$$

Subsequently, we evaluate $\frac{\partial}{\partial \eta}\left(\frac{\partial W}{\partial \zeta}(\eta,\eta)\right)$ at $\eta^* = \frac{D}{2}$.

$$\frac{\partial}{\partial \eta}\left(\frac{\partial W}{\partial \zeta}(\eta,\eta)\right)(\frac{D}{2}) = -\frac{\delta}{\tau^2} - \frac{\varepsilon D}{2\tau^2}a_2(\frac{D}{2})\frac{\frac{D}{2}-D}{\tau^2}R_2(\frac{D}{2}) + \frac{\varepsilon D}{2\tau^2}a_2(\frac{D}{2})\frac{\partial R_2}{\partial \eta}(\frac{D}{2}). \tag{SI3.16}$$

Evaluating $\frac{\partial R_2}{\partial \eta}(\eta)$ at $\eta^* = \frac{D}{2}$, we get

$$\begin{aligned}\frac{\partial R_2}{\partial \eta}\left(\frac{D}{2}\right) &= -\frac{\delta}{\varepsilon}\frac{1}{a_2(\frac{D}{2})}\frac{\frac{D}{2\tau^2}a_2(\frac{D}{2})F^*(\frac{D}{2})}{a_2(\frac{D}{2})F^*(\frac{D}{2})+\rho} \\ &= -\frac{D\delta}{2\varepsilon\tau^2}\frac{F^*(\frac{D}{2})}{a_2(\frac{D}{2})F^*(\frac{D}{2})+\rho} \\ &= -\frac{D\delta}{2\varepsilon\tau^2}(-\frac{\delta}{2\varepsilon R_{max}a_2(\frac{D}{2})^2} + \frac{1}{a_2(\frac{D}{2})})\end{aligned} \tag{SI3.17}$$

because

$$\begin{aligned}F^*\left(\frac{D}{2}\right) &= \frac{-2\rho\delta a_1(\frac{D}{2}) + \varepsilon\rho R_{max}a_1(\frac{D}{2})^2 + \rho\varepsilon R_{max}a_1(\frac{D}{2})^2}{2\delta a_1(\frac{D}{2})^2} \\ &= \frac{-\rho\delta + \varepsilon\rho R_{max}a_1(\frac{D}{2})}{\delta a_1(\frac{D}{2})} \\ &= \frac{-\rho}{a_1(\frac{D}{2})} + \frac{\varepsilon\rho R_{max}}{\delta}\end{aligned} \tag{SI3.18}$$

Replacing eq. (SI3.17) in eq. (SI3.16), it yields

$$\begin{aligned}
\frac{\partial}{\partial\eta}\left(\frac{\partial W}{\partial\zeta}(\eta,\eta)\right)\left(\frac{D}{2}\right) &= -\frac{\delta}{\tau^2}+\frac{\delta D^2}{4\tau^4}+\frac{\varepsilon D}{2\tau^2}a_2(\frac{D}{2})(-\frac{D\delta}{2\varepsilon\tau^2}(-\frac{\delta}{\varepsilon R_{max}a_2(\frac{D}{2})^2}+\frac{1}{a_2(\frac{D}{2})})) \\
&= -\frac{\delta}{\tau^2}+\frac{\delta D^2}{4\tau^4}+\frac{\delta D^2}{4\tau^4}a_2(\frac{D}{2})(\frac{\delta}{\varepsilon R_{max}a_2(\frac{D}{2})^2}-\frac{1}{a_2(\frac{D}{2})})) \\
&= -\frac{\delta}{\tau^2}+\frac{\delta^2 D^2}{4\varepsilon R_{max}\tau^4}\frac{1}{a_2(\frac{D}{2})} \\
&= -\frac{\delta}{\tau^2}+\frac{\delta^2 D^2}{4\varepsilon R_{max}\tau^4 A}e^{\frac{D^2}{8\tau^2}} \qquad \text{(SI3.19)}
\end{aligned}$$

Because $\frac{\partial}{\partial\eta}\left(\frac{\partial W}{\partial\zeta}(\eta,\eta)\right)(\frac{D}{2})$ is an increasing function of $D$, there exists $\bar{D}$ such that $\frac{\partial}{\partial\eta}\left(\frac{\partial W}{\partial\zeta}(\frac{D}{2},\frac{D}{2})\right) < 0$ for any $D < \bar{D}$ and $\frac{\partial}{\partial\eta}\left(\frac{\partial W}{\partial\zeta}(\frac{D}{2},\frac{D}{2})\right) > 0$ for any $D > \bar{D}$. Thus, $\bar{D}$ is the unique solution of the equation $\frac{\partial}{\partial\eta}\left(\frac{\partial W}{\partial\zeta}(\eta,\eta)\right)(\frac{D}{2}) = 0$ which we rewrite using equation (SI3.19):

$$\begin{aligned}
0 &= -\frac{\delta}{\tau^2}+\frac{\delta^2 D^2}{4\varepsilon R_{max}\tau^4 A}e^{\frac{D^2}{8\tau^2}} \qquad \text{(SI3.20)} \\
&= -\frac{1}{\tau^2}+\frac{\delta D^2}{4\varepsilon R_{max}\tau^4 A}e^{\frac{D^2}{8\tau^2}} \qquad \text{(SI3.21)}
\end{aligned}$$

Equation (SI3.21) is equivalent to

$$D^2 e^{\frac{D^2}{8\tau^2}} = \frac{4\varepsilon R_{max}\tau^2 A}{\delta} \qquad \text{(SI3.22)}$$

Therefore, for any $R_{max}$ larger than $\frac{\delta D^2}{4\varepsilon\tau^2 A}e^{\frac{D^2}{8\tau^2}}$, the evolutionary equilibrium $\eta^* = \frac{D}{2}$ is stable and thus an evolutionary attractor.

In conclusion, diversification occurs at the evolutionary equilibrium $\eta^* = \frac{D}{2}$ when the two following conditions are met:

$$\begin{aligned}
2\tau &< D \qquad \text{(SI3.23)} \\
\frac{\delta D^2}{4\varepsilon\tau^2 A}e^{\frac{D^2}{8\tau^2}} &< R_{max}. \qquad \text{(SI3.24)}
\end{aligned}$$

The second condition can be rewritten in terms of the productivity of the system, i.e. $P = R_{1max} + R_{2max} = 2R_{max}$:

$$\frac{\delta D^2}{2\varepsilon\tau^2 A}e^{\frac{D^2}{8\tau^2}} < P. \qquad \text{(SI3.25)}$$

$$\frac{\partial^2 L(\eta_k\prime, \eta)}{\partial \eta_k\prime^2} = \frac{\varepsilon}{2\delta} \sum_{i=1}^{n} A \frac{((\theta_i - \eta_k\prime) - \tau^2)}{\tau^4} e^{\frac{-(\theta_i - \eta_k\prime)^2}{2\tau^2}} R_i^*(\eta) \tag{SI2.4}$$

When equation (SI2.4) is positive, selection is disruptive and diversification (or evolutionary branching, according to adaptive dynamics theory) occurs.

**Asexual lineage** To enable direct analytical comparison between the reproductive modes, we also use the lifetime reproductive output as a measure of invasion fitness to model the evolutionary dynamics of the asexual lineage. In this case, the lifetime reproductive output of a rare mutant $\eta_j\prime$ at ecological equilibrium when resident ecomorphs have traits $\eta = (\eta_1, \ldots, \eta_m)$ is given by the product of the average lifetime of an individual, $1/\delta$, and its reproduction rate, thus:

$$L(\eta_j\prime, \eta) = \frac{\varepsilon}{\delta} \sum_{i=1}^{n} a_i(\eta_j\prime) R_i^*(\eta) \tag{SI2.5}$$

Hence, the dynamics of the feeding niche trait of the $j$th ecomorph population follows:

$$\frac{d\eta_j}{dt} = \sigma_a \frac{\partial L(\eta_j\prime, \eta)}{\partial \eta_j\prime} \tag{SI2.6}$$

Because we use $L$ as proxy for fitness, the fitness gradient $\frac{\partial s(\eta_j\prime, \eta)}{\partial \eta_{j'}}$ found in eq. 6 in Methods is replaced by $\frac{\partial L(\eta_j\prime, \eta)}{\partial \eta_{j'}}$.

From eq. SI2.5, we have

$$\frac{\partial L(\eta_j\prime, \eta)}{\partial \eta_j\prime} = \frac{\varepsilon}{\delta} \sum_{i=1}^{n} A \frac{(\theta_i - \eta_j\prime)}{\tau^2} e^{\frac{-(\theta_i - \eta_j\prime)^2}{2\tau^2}} R_i^*(\eta) \tag{SI2.7}$$

Directional selection halts where the selection gradient vanishes, in other words when equation (SI2.7) equals 0. At this point, selection can be stabilizing if this trait value corresponds to a fitness maximum, or disruptive if it corresponds to a fitness minimum. The curvature of the fitness function therefore determines whether evolution halts or an ecomorph splits into two different ecomorphs. Hence, the second derivative of the fitness function with respect to $\eta_j\prime$ evaluated at the ecological equilibrium when resident ecomorphs have traits $\eta$ allows us to determine whether a diversification event occurs:

$$\frac{\partial^2 L(\eta_j\prime, \eta)}{\partial \eta_j\prime^2} = \frac{\varepsilon}{\delta} \sum_{i=1}^{n} A \frac{((\theta_i - \eta_j\prime) - \tau^2)}{\tau^4} e^{\frac{-(\theta_i - \eta_j\prime)^2}{2\tau^2}} R_i^*(\eta) \tag{SI2.8}$$

When equation (SI2.8) is positive, selection is disruptive and diversification (or evolutionary branching, according to adaptive dynamics theory) occurs.

**References** /

1. J. A. J. Metz, C. G. F. de Kovel, The canonical equation of adaptive dynamics for Mendelian diploids and haplodiploids. Interface Focus 3 (2013).

### Supplementary Information 4. Analytical proof that the density of an asexual population is higher than the density of a sexual population at ecological equilibrium

Here we show that the density of an asexual population ($N_a$) is always higher than the density of a sexual population ($N_s$) at ecological equilibrium. To do so, we first derive analytical expressions for the population and resource densities at ecological equilibrium for an asexual and a sexual population. We then compare the population densities of these populations.

**Ecological equilibrium of an asexual population** At ecological equilibrium, the population density, $N_a$, and resources, $R_1$ and $R_2$, satisfy the following equations

$$\begin{aligned} 0 &= \varepsilon(a_1(\eta)R_1 + a_2(\eta)R_2) - \delta && \text{(SI4.1)} \\ 0 &= \rho(R_{max} - R_1) - a_1(\eta)R_1N_a && \text{(SI4.2)} \\ 0 &= \rho(R_{max} - R_2) - a_2(\eta)R_2N_a && \text{(SI4.3)} \end{aligned}$$

Equations (SI4.2) and (SI4.3) give us

$$R_1 = \frac{\rho R_{max}}{\rho + a_1(\eta)N_a} \text{ and } R_2 = \frac{\rho R_{max}}{\rho + a_2(\eta)N_a} \quad \text{(SI4.4)}$$

By substituting $R_1$ and $R_2$ in equation (SI4.1), we obtain an equation for $N_a$:

$$0 = \varepsilon(a_1(\eta)\frac{\rho R_{max}}{\rho + a_1(\eta)N_a} + a_2(\eta)\frac{\rho R_{max}}{\rho + a_2(\eta)N_a}) - \delta$$

which is equivalent to the quadratic equation

$$0 = \alpha_a N_a^2 + \beta_a N_a + \gamma_a \quad \text{(SI4.5)}$$

where

$$\begin{aligned} \alpha_a &= \delta a_1(\eta)a_2(\eta) && \text{(SI4.6)} \\ \beta_a &= \rho\delta a_1(\eta) + \rho\delta a_2(\eta) - 2\varepsilon\rho R_{max}a_1(\eta)a_2(\eta) && \text{(SI4.7)} \\ \gamma_a &= \rho^2\delta - \varepsilon\rho^2 R_{max}(a_1(\eta) + a_2(\eta)) && \text{(SI4.8)} \end{aligned}$$

The two solutions of equation (SI4.5) are

$$N_a^*(\eta) = \frac{-\beta_a \pm \sqrt{\Delta_a}}{2\alpha_a}$$

where $\Delta_a = \beta_a^2 - 4\alpha_a\gamma_a$. Only when $\sqrt{\Delta_a}$ is added to $-\beta_a$ the population density is positive, therefore, the only biologically relevant solution is

$$N_a^*(\eta) = \frac{-\beta_a + \sqrt{\Delta_a}}{2\alpha_a} \quad \text{(SI4.9)}$$

Replacing $N_a^*$ in equations (SI4.4), we obtain:

$$R_1^*(\eta) = \frac{2a_2(\eta)R_{max}\delta\rho}{\sqrt{\rho^2(4b^2+\delta^2(a_2(\eta)-a_1(\eta))^2)}+\rho(2b+\delta(a_2(\eta)-a_1(\eta)))} \quad \text{(SI4.10)}$$

$$R_2^*(\eta) = \frac{2a_1(\eta)R_{max}\delta\rho}{\sqrt{\rho^2(4b^2+\delta^2(a_2(\eta)-a_1(\eta))^2)}+\rho(2b-\delta(a_2(\eta)-a_1(\eta)))}, \quad \text{(SI4.11)}$$

where $b = a_1(\eta)a_2(\eta)\varepsilon R_{max}$.

**Ecological equilibrium of a sexual population** At ecological equilibrium, the density of females, $F$, males, $M$, and resources, $R_1$ and $R_2$, satisfy the following equations

$$0 = \frac{1}{2}\varepsilon(a_1(\eta)R_1 + a_2(\eta)R_2) - \delta \quad \text{(SI4.12)}$$

$$0 = \frac{1}{2}\varepsilon(a_1(\eta)R_1 + a_2(\eta)R_2)F - \delta M \quad \text{(SI4.13)}$$

$$0 = \rho(R_{max} - R_1) - a_1(\eta)R_1(F+M) \quad \text{(SI4.14)}$$

$$0 = \rho(R_{max} - R_2) - a_2(\eta)R_2(F+M) \quad \text{(SI4.15)}$$

Because the primary sex ratio is 1/2, and the mortality and feeding rate is equal for females and males, then $M = F$. Thus the system can be reduced to a system of three equations:

$$0 = \frac{1}{2}\varepsilon(a_1(\eta)R_1 + a_2(\eta)R_2) - \delta \quad \text{(SI4.16)}$$

$$0 = \rho(R_{max} - R_1) - a_1(\eta)R_1(2F) \quad \text{(SI4.17)}$$

$$0 = \rho(R_{max} - R_2) - a_2(\eta)R_2(2F) \quad \text{(SI4.18)}$$

Equations (SI4.17) and (SI4.18) give us

$$R_1 = \frac{\rho R_{max}}{\rho + a_1(\eta)(2F)} \text{ and } R_2 = \frac{\rho R_{max}}{\rho + a_2(\eta)(2F)} \quad \text{(SI4.19)}$$

By substituting $R_1$ and $R_2$ in equation (SI4.16), we obtain an equation for $F$:

$$0 = \frac{1}{2}\varepsilon(a_1(\eta)\frac{\rho R_{max}}{\rho + a_1(\eta)(2F)} + a_2(\eta)\frac{\rho R_{max}}{\rho + a_2(\eta)(2F)}) - \delta$$

which is equivalent to the quadratic equation

$$0 = \alpha_s(2F)^2 + \beta_s(2F) + \gamma_s \quad \text{(SI4.20)}$$

where

$$\alpha_s = \delta a_1(\eta)a_2(\eta) \quad \text{(SI4.21)}$$

$$\beta_s = \rho\delta a_1(\eta) + \rho\delta a_2(\eta) - \varepsilon\rho R_{max}a_1(\eta)a_2(\eta) \quad \text{(SI4.22)}$$

$$\gamma_s = \rho^2\delta - \frac{1}{2}\varepsilon\rho^2 R_{max}(a_1(\eta) + a_2(\eta)) \quad \text{(SI4.23)}$$

Note that $\alpha_s = \alpha_a$ but $\beta_s > \beta_a$ and $\gamma_s > \gamma_a$.
The two solutions of equation (SI4.20) are

$$(2F)^*(\eta) = \frac{-\beta_s \pm \sqrt{\Delta_s}}{2\alpha_s}$$

where $\Delta_s = \beta_s^2 - 4\alpha_s\gamma_s$. Only when $\sqrt{\Delta_s}$ is added to $-\beta_s$ the population density is positive, therefore, the only biologically relevant solution is

$$(2F)^*(\eta) = \frac{-\beta_s + \sqrt{\Delta_s}}{2\alpha_s} \tag{SI4.24}$$

Replacing $F^*$ in equations (SI4.19), we obtain:

$$R_1^*(\eta) = \frac{2a_2(\eta)R_{max}\delta\rho}{\sqrt{\rho^2\left(b^2 + \delta^2(a_2(\eta) - a_1(\eta))^2\right)} + \rho(b + \delta(a_2(\eta) - a_1(\eta)))} \tag{SI4.25}$$

$$R_2^*(\eta) = \frac{2a_1(\eta)R_{max}\delta\rho}{\sqrt{\rho^2\left(b^2 + \delta^2(a_2(\eta) - a_1(\eta))^2\right)} + \rho(b - \delta(a_2(\eta) - a_1(\eta)))}, \tag{SI4.26}$$

where $b = a_1(\eta)a_2(\eta)\varepsilon R_{max}$.
Note that the analytical expressions obtained for $R_1^*$ and $R_2^*$ in an asexual population (eq. SI4.10 and eq. SI4.11) are different to the expressions obtained for $R_1^*$ and $R_2^*$ in a sexual population (eq. SI4.25 and eq. SI4.26). In Supplementary Information 5, we demonstrate that due to these differences, selection is always stronger in an asexual population than in a sexual population.

**Comparing the density of an asexual and a sexual population** In this paragraph, we proof that an asexual population has a higher density than a sexual population. Let us define

$$\beta(t) = \rho\delta a_1(\eta) + \rho\delta a_2(\eta) - t\varepsilon\rho R_{max}a_1(\eta)a_2(\eta) \tag{SI4.27}$$

$$\gamma(t) = \rho^2\delta - t\frac{1}{2}\varepsilon\rho^2 R_{max}(a_1(\eta) + a_2(\eta)). \tag{SI4.28}$$

From eq.SI4.9, we have

$$N_a^*(\eta) = \frac{-\beta(2) + \sqrt{\beta(2)^2 - 4\alpha\gamma(2)}}{2\alpha} \tag{SI4.29}$$

where

$$\alpha = \delta a_1(\eta)a_2(\eta) \tag{SI4.30}$$

From eq. SI4.24, we have

$$N_s^*(\eta) = (2F)^*(\eta) = \frac{-\beta(1) + \sqrt{\beta(1)^2 - 4\alpha\gamma(1)}}{2\alpha}. \tag{SI4.31}$$

We want to show that $N_a^*(\eta) > N_s^*(\eta)$. Let us define the function

$$r(t) = \frac{-\beta(t) + \sqrt{\beta(t)^2 - 4\alpha\gamma(t)}}{2\alpha}, \tag{SI4.32}$$

From equations (SI4.29) and (SI4.31), we can show that $N_a^*(\eta) = r(2)$ and $N_s^*(\eta) = r(1)$. Therefore, our claim is proven if we show that $r$ is an increasing function of $t$. Let us calculate its derivative.

$$\frac{dr}{dt} = \frac{1}{2\alpha}\left(-\frac{d\beta}{dt} + \frac{\beta(t)\frac{d\beta}{dt} - 2\alpha\frac{d\gamma}{dt}}{\sqrt{\beta(t)^2 - 4\alpha\gamma(t)}}\right) \tag{SI4.33}$$

$$= \frac{1}{2\alpha}\left(\varepsilon\rho R_{max}a_1(\eta)a_2(\eta) + \frac{\varepsilon^2\rho^2 R_{max}^2 a_1(\eta)^2 a_2(\eta)^2 t}{\sqrt{\beta(t)^2 - 4\alpha\gamma(t)}}\right) \tag{SI4.34}$$

$$> 0. \tag{SI4.35}$$

### Supplementary Information 5. Analytical investigation of the strength of selection in populations with different reproductive modes

Here, we analytically investigate whether the reproductive mode has an effect on the strength of selection of a population colonizing an environment with two different food resources, $R_1$ and $R_2$. We restrict our analytical investigation to the simplest case, in which $R_{1max} = R_{2max} = R_{max}$. The optimal niche trait values to feed on resource 1 and resource 2 are $\theta_1$ and $\theta_2$, respectively, and their distance in the trait axis equals $D$. Without a loss of generality, we assume that $\theta_1 = 0$ and $\theta_2 = D$. Following the adaptive dynamics framework, we assume that evolution occurs much more slowly than the ecological dynamics. This separation of the ecological and evolutionary timescales enables us to approximate the ecological dynamics by assuming that they are always in a quasi-steady state determined by the ecomorph population with trait value $\eta$. Using the analytical expressions for the population and resource densities at ecological equilibrium for an asexual and a sexual population derived in SI4, we investigate the selection gradient of a rare mutant in the environment set by the population with a trait value $\eta$ for each reproductive mode. Finally, we compare the selection gradient of both populations (sexual and asexual), and demonstrate that selection is stronger in an asexual than in a sexual population.

**Selection gradient of an asexual population** Using eq. (SI2.1), the fitness gradient, calculated using the lifetime reproductive output, of a rare mutant $\eta\prime$ at ecological equilibrium when the resident population has trait $\eta$ is given by

$$\frac{\partial L(\eta\prime,\eta)}{\partial \eta\prime} = \frac{\varepsilon}{\delta} A \left( \frac{(-\eta\prime)}{\tau^2} e^{\frac{-(-\eta\prime)^2}{2\tau^2}} R_1^*(\eta) + \frac{(D-\eta\prime)}{\tau^2} e^{\frac{-(D-\eta\prime)^2}{2\tau^2}} R_2^*(\eta) \right) \tag{SI5.1}$$

By substituting equations (SI4.10) and (SI4.11) in (SI5.1) and after some algebraic manipulation, we obtain the following expression for the selection gradient:

$$S_{asexual} = \frac{\partial L(\eta\prime,\eta)}{\partial \eta\prime} = \frac{2\varepsilon R_{max} \rho A^2}{\tau^2} e^{\frac{-(2\eta^2 - 2D\eta + D^2)}{2\tau^2}} C_{asexual}, \tag{SI5.2}$$

where

$$C_{asexual} = \frac{-\eta}{\left(\sqrt{\rho^2(4b^2 + \delta^2(a_2(\eta) - a_1(\eta))^2)} + \rho(2b + \delta(a_2(\eta) - a_1(\eta)))\right)} + \frac{D-\eta}{\sqrt{\rho^2(4b^2 + \delta^2(a_2(\eta) - a_1(\eta))^2)} + \rho(2b - \delta(a_2(\eta) - a_1(\eta)))} \tag{SI5.3}$$

**Selection gradient of a sexual population** Using equation (SI2.3), the fitness gradient, calculated using the lifetime reproductive output, of a rare mutant $\eta\prime$ at ecological equilibrium when the resident population has trait $\eta$ is given by

$$\frac{\partial L(\eta\prime,\eta)}{\partial \eta\prime} = \frac{\varepsilon}{2\delta}A\left(\frac{(-\eta\prime)}{\tau^2}e^{\frac{-(-\eta\prime)^2}{2\tau^2}}R_1^*(\eta) + \frac{(D-\eta\prime)}{\tau^2}e^{\frac{-(D-\eta\prime)^2}{2\tau^2}}R_2^*(\eta)\right) \tag{SI5.4}$$

By substituting equations (SI4.25) and (SI4.26) in (SI5.4) and after some algebraic manipulation, we obtain the following expression for the selection gradient:

$$S_{sexual} = \frac{\partial L(\eta\prime,\eta)}{\partial \eta\prime} = \frac{\varepsilon R_{max}\rho A^2}{\tau^2}e^{\frac{-(2\eta^2-2D\eta+D^2)}{2\tau^2}}C_{sexual}, \tag{SI5.5}$$

where

$$C_{sexual} = \frac{-\eta}{\left(\sqrt{\rho^2(b^2+\delta^2(a_2(\eta)-a_1(\eta))^2)}+\rho(b+\delta(a_2(\eta)-a_1(\eta)))\right)} + \frac{D-\eta}{\sqrt{\rho^2(b^2+\delta^2(a_2(\eta)-a_1(\eta))^2)}+\rho(b-\delta(a_2(\eta)-a_1(\eta)))} \tag{SI5.6}$$

**Comparing the selection gradient of an asexual and a sexual population** In this paragraph we prove that the selection pressure driving the trait to the value where selection is disruptive, and thus where diversification occurs, is always smaller in a sexual population than in an asexual population. Diversification occurs when $\eta = \frac{D}{2}$ is an attractor of the evolutionary dynamics because this evolutionary equilibrium corresponds to a local minimum of the fitness landscape (see SI3). When $\eta = \frac{D}{2}$ is an attractor for both sexual and asexual populations, there exists $\zeta > 0$ such that $S_{sexual}, S_{asexual} > 0$ for $\zeta < \eta < \frac{D}{2}$ and $S_{sexual}, S_{asexual} < 0$ for $\frac{D}{2} < \eta < \zeta$. Hence, we want to show that $0 < S_{sexual} < S_{asexual}$ for $\zeta < \eta < \frac{D}{2}$ and $0 > S_{sexual} > S_{asexual}$ for $\frac{D}{2} < \eta < \zeta$. From equations (SI5.2) and (SI5.5), this is equivalent to show that $0 < C_{sexual} < C_{asexual}$ for $\zeta < \eta < \frac{D}{2}$ and $0 > C_{sexual} > C_{asexual}$ for $\frac{D}{2} < \eta < \zeta$.

Let us define the function

$$z(t) = \frac{-2t\eta\sqrt{t^2\phi^2+\mu^2} - 2t^2\eta\phi + tD\sqrt{t^2\phi^2+\mu^2} + t^2D\phi + tD\mu}{2t^2\phi^2 + 2t\phi\sqrt{t^2\phi^2+\mu^2}} \tag{SI5.7}$$

$$= \frac{(D-2\eta)(\sqrt{t^2\phi^2+\mu^2}+t\phi)+D\mu}{2\phi(t\phi+\sqrt{t^2\phi^2+\mu^2})}, \tag{SI5.8}$$

where $\phi = \rho\varepsilon R_{max}a_2(\eta)a_1(\eta)$ and $\mu = \rho\delta(a_2(\eta)-a_1(\eta))$. Using equations (SI5.3) and (SI5.6), we can show that $z(1) = C_{sexual}$ and $z(2) = C_{asexual}$. Therefore our claim is proven if $z$ is an increasing function in $t$ when $\eta < \frac{D}{2}$ and it is a decreasing function when $\eta > \frac{D}{2}$. We then calculate the derivative of the function $z$. First we derive the numerator (that we name $J$) of $z$ and then the denominator (that we name $K$) of $z$:

$$\frac{dJ}{dt} = (D-2\eta)\left(\frac{2t\phi^2}{2\sqrt{t^2\phi^2+\mu^2}}+\phi\right) \tag{SI5.9}$$

$$= \phi(D-2\eta)\left(\frac{t\phi}{\sqrt{t^2\phi^2+\mu^2}}+1\right) \tag{SI5.10}$$

$$\frac{dK}{dt} = 2\phi^2(1+\frac{t\phi}{\sqrt{t^2\phi^2+\mu^2}}) \tag{SI5.11}$$

The sign of the derivative $\frac{dz}{dt} > 0$ is determined by the sign of $\frac{dJ}{dt}K - \frac{dK}{dt}J$ which is given by

$$\frac{dJ}{dt}K - \frac{dK}{dt}J = -2\phi^2D\mu(1+\frac{t\phi}{\sqrt{t^2\phi^2+\mu^2}}). \tag{SI5.12}$$

Since $\phi$ and $D$ are positive, the sign of the $\frac{dz}{dt} > 0$ is determined by the term $\mu$. The sign of $\mu$ is positive when $a_2(\eta) > a_1(\eta)$, which occurs when $\eta > \frac{D}{2}$. Conversely, $\mu$ is negative when $\eta < \frac{D}{2}$, which concludes the proof.

### Supplementary Information 6. Individual-based model details

In the deterministic model, the dynamics of the system describe the change of consumer densities. In contrast, in the IBM, individual consumers are discrete entities. We thus consider $l$ females ($k = 1, \dots, l$) and $r$ males ($r = 1, \dots, s$) in the case of the sexual lineage, and $m$ clonal individuals ($j = 1, \dots, m$) in the case of the asexual lineage.

In the IBM, the change in density of the $i$th resource $R_i$ in a time step $\Delta t$ equals

$$\Delta R_i = \left[\rho(R_{i\,\max} - R_i) - \left(\sum_{k=1}^{l} a_i(\eta_k)R_i + \sum_{r=1}^{s} a_i(\eta_r)R_i\right)/V\right]\Delta t \tag{SI6.1}$$

in the case of the sexual lineage, and

$$\Delta R_i = \left[\rho(R_{i\,\max} - R_i) - \left(\sum_{j=1}^{m} a_i(\eta_j)R_i\right)/V\right]\Delta t \tag{SI6.2}$$

in the case of the asexual lineage. In eq. SI6.1 and eq. SI6.2, $V$ is the volume of the system, which divides the the intake of resource $i$ by all individuals. This way, we can calculate the change in resource density due to consumption.

We assume that all reproducing individuals (sexual females and clonal individuals) have a reproductive buffer $B$, which increases over time following

$$\Delta B = \left[\varepsilon \sum_{i=1}^{n} a_i(\eta_j)R_i\right]\Delta t \tag{SI6.3}$$

The reproductive buffer can take values between 0 and 1. The probability that a reproducing individual (sexual female or clonal individual) actually reproduces depends on the value of the buffer, such that its probability to reproduce equals this value. If an individual reproduces, it produces one individual offspring and its reproductive buffer is set to 0. Although males have the same feeding rate than sexual females (as shown by eq. SI6.2), the food consumed is not used for reproduction. While it may be biologically unrealistic (especially if males provide parental care), this assumption ensures that the cost of producing male offspring in the IBM is equal to the analogous cost in the adaptive dynamics model.

In the asexual lineage, a clonal individual produces one offspring that inherits the total maternal genetic material (without recombination). In the sexual lineage, a sexual female and a sexual male produce one offspring that receives in each locus one allele from the father and one from the mother with probability, $p$, or both alleles from the mother with probability, $(1-p)$. In each birth event, each E-locus has a probability $\mu_E$ to mutate, which is independent of the mutation probability of other loci. In such case, the offspring allele equals the allele of the parent + $\gamma$, with $\gamma$ normally distributed with a mean of zero and standard deviation $\sigma$. Additionally, in the sexual lineage, with probability $\mu_A$, a mutation occurs independently in each A-locus, reversing the sign of the allele value (as modeled in ref (1)). To determine which offspring mutate in which allele, we draw for each allele for each newborn a random number from a uniform distribution on the interval [0,1]. If this number is smaller than the mutation rate $\mu_E$, in the case of an E-locus, or $\mu_A$, in the case of a A-locus, this allele mutates.

To describe mating in the sexual lineage, we follow Dieckmann and Doebeli (1). We assume that mating depends on female preference. The probability that a female with assortative mating trait $\omega_k$ and feeding niche trait $\eta_k$ will mate with a male with niche trait $\eta_r$ is described by the following gaussian function:

$$P(\omega_k, \eta_k, \eta_r) = \begin{cases} \dfrac{{\omega_k}^2}{\nu_a\sqrt{2\pi}} e^{-\frac{1}{2}\left(\frac{{\omega_k}^2(\eta_k-\eta_r)}{\nu_a}\right)} & \omega_k > 0 \\ 1 & \omega_k = 0 \\ 1 - \dfrac{{\omega_k}^2}{\nu_s\sqrt{2\pi}} e^{-\frac{1}{2}\left(\frac{{\omega_k}^2(\eta_k-\eta_r)}{\nu_s}\right)} & \omega_k < 0 \end{cases} \tag{SI6.4}$$

In this equation, $\nu_a$ and $\nu_s$ are scaling parameters determining the slope of the mate-choice function. To avoid a bias against rare phenotypes, mating probabilities are normalized, such that the sum of mating probabilities over all potential partners is 1 for each female, irrespective of her own phenotype. This implies that there is no cost to a female for assortative mating.

For each time step $\Delta t$, we first calculate the food consumption by each individual and update their reproductive buffer. Next, we update the food resource density using eq. SI6.1 or SI6.2 for the sexual and asexual lineage, respectively. Then, reproduction takes place. Finally, we remove individuals that have died. To determine which individuals die at each time step, we draw for each individual a random number from a uniform distribution on the interval [0,1]. If this number is smaller than the mortality rate $\delta$, this individual dies.

To calculate the level of reproductive isolation in the sexual lineage, we count the number of matings between conspecifics and heterospecifics during the last 1000 time steps of a simulation. A mating is considered to occur between conspecific if the feeding niche trait of the two individuals fall within the range of the same discrete cluster (see figure S3B), and between heterospecific if the feeding niche trait of the two individuals fall in the range of different discrete clusters. Based on the number of conspecific and heterospecific matings, we use the statistic proposed in ref (2) to compute the level of reproductive isolation.

We simulate the diversification process of an asexual (figure 3A) and a sexual (figure 3B) lineage independently, and when they cooccur in the same environment (figure 3E). In the latter case, both founder populations are seeded at the beginning of the simulation in the extremes of the trait space, ensuring that they have equal access to the trait space. Because the food resources are shared by both lineages, their dynamics thus follow:

$$\Delta R_i = \left[\rho(R_{i\,\max} - R_i) - \left(\sum_{k=1}^{l} a_i(\eta_k)R_i + \sum_{r=1}^{s} a_i(\eta_r)R_i + \sum_{j=1}^{m} a_i(\eta_j)R_i\right)/V\right]\Delta t$$

(SI2.5)

We also run 100 simulations of a sexual lineage when $p = (0,0.1,0.2,\dots,1)$ and of an asexual lineage as reference (figure 4), and save the first point in time when all food resources have been utilized (we define this point as the first time when all food resources have a density smaller than half of their carrying capacity, i.e. $R_i < \frac{1}{2}R_{imax}$).

*Parameterization of the IBM:* The model is implemented based on the 'Model assumptions' describe in the Methods. Therefore, the demographic and environmental parameters used in the adaptive dynamics model (table S1) are also in the IBM. Additionally, in the IBM, it is necessary to select an appropriate habitat size to minimize the effects of demographic stochasticity. To select this parameter value, we perform simulations of the IBM of the asexual lineage with different habitat size (volume) assuming no mutation (only ecological dynamics) (see figure S9). Based on the coefficient of variation of individual density and the computational time of these simulations, we select a habitat size of 1000. We use the values of genetic parameters that Dieckmann and Doebeli previously used in the IBM that form the basis of our model (1). However, we performed robustness test with different values of genetic parameters. Specifically, we vary the mutation rate ($10^{-3}$, $10^{-4}$) and number of E-loci (5, 10, 15, 20). The results are presented in figure S8.

Table S2. Parameters of the IBM

| Variable or parameter | Symbol | Value | Units |
|---|---|---|---|
| Variables | | | |
| Feeding niche trait | $\eta$ | Evolving trait | - |
| Mating trait (only in sexually reproducing individuals) | $\omega$ | Evolving trait | - |
| Number of clonal individuals | $m$ | - | $L^{-1}$ |
| Number of females | $l$ | - | $L^{-1}$ |
| Number of males | $r$ | - | $L^{-1}$ |
| Food resource density | $R_i$ | - | g $L^{-1}$ |
| Environmental parameters | | | |
| Habitat size | $V$ | 1000 | L |
| Genetic parameters | | | |
| Number of E-loci | - | 10 (by default) varied in fig. S8 | - |
| Number of A-loci** | - | 5 | - |
| Mutation rate of E-loci | $\mu_E$ | $10^{-3}$ (by default) varied in fig. S8 | - |
| Mutation rate of A-loci** | $\mu_A$ | $10^{-3}$ | - |
| Standard deviation of the mutation distribution | $\sigma$ | 0.01 | - |
| Probability of receiving one allele from the mother and one allele from the father in each locus | $p$ | 1 (by default) varied in fig. 5 | - |

** Parameter applies only to sexual population(s)

*Initialization of IBM simulations:* All IBM simulations were initialized with one (in the case of figure 3A, 3B and 4) or two populations (in the case of figure S6A) composed of 40 individuals with identical trait values (20 females and 20 males in a sexual population). In the case that only one ancestral population occurs, the feeding niche trait of all individuals is 0.8. In the case that both a sexual and an asexual population co-occur, the feeding niche trait of the individuals in the sexual population is 10.2 and in the asexual population is 0.8 (the value of each allele in the E-locus set equals the trait value divided by the double -because of diploidy- of the number of E-locus, such that their sum is the feeding niche trait). These trait values ensure that both populations are located in the extremes of the trait space and thus that they have equal access to the trait space. In this way, their process of trait expansion is interfered by each other only when all niches have been occupied. The ancestral sexual populations are initialized assigning to each individual a mating trait $\omega = 0$ (i.e. individuals carry an equal number of 1 and -1 alleles in the A-locus set such that their average equals 0), hence they mate randomly.

*Investigating the stability of the attractor of a sexual lineage*

In addition, we examine the stability of the attractor of a sexual lineage (final state in figure 3B), by simulating different types of disturbances (figure S7) as well as seeding a simulation with multiple initial populations (figure S10). A detailed discussion of these results can be found in SI7.

### Supplementary Information 7. Investigation of stability of the final state of a sexual lineage

We investigate the stability of the final state of the sexual lineage by simulating diverse disturbance events. First, we simulate an event in which half of the individuals die randomly at a specific point in time (figure S7A, time 1000). Second, we simulate an event in which all individuals with a feeding niche trait larger than 7.5 die, in this case 4 populations get extinct (figure S7B). Third, we simulate a sustain perturbation, in which the mortality rate increases gradually from 0.02 to 0.08 in steps of 0.001 per unit of time (figure S7C). In all cases, the lineage does not collapse into a single population and all populations recover back rapidly from the temporary perturbation (in the case of the first and second disturbance) or persists at low abundance despite the high mortality.

We also simulate an extinction event in an environment in which the sexual and the asexual lineages co-occur (figure S6B). This event leads to the extinction of all individuals with trait values between 2.5 and 4.5, that is the individuals in the boundary between the lineages with alternative reproductive modes. After the event, two possible alternative scenarios are observed: In the first (observed in 6 out of 10 simulations), each lineage reoccupies its original niche space, such that food resource 3 and 4 are again used by the asexual and sexual lineage, respectively. In the second (observed in 4 out of 10 simulations), the sexual lineage rapidly reoccupies the available niche that was previously occupying (near the optimal of resource 4) and additionally occupies the vacant niche left by the extinct asexual population (near the optimum of resource 3).

Additionally, we simulate a sexual lineage seeding the simulation with discrete populations composed of individuals whose feeding niche trait equals each of the optima to feed on the resources and mating trait equals 1, such that assortative mating is maximal (figure S10). After short time these populations mix up and a hybrid swarm, as observed before time 1000000 in figure 3B, quickly emerges. Later, discrete populations emerge again but they are a combination of generalists and specialists, in the exact way that is observed at the end of the simulation in figure 3B. These results suggest that the state observed at the end of the simulation in figure 3B is an attractor with large stability.

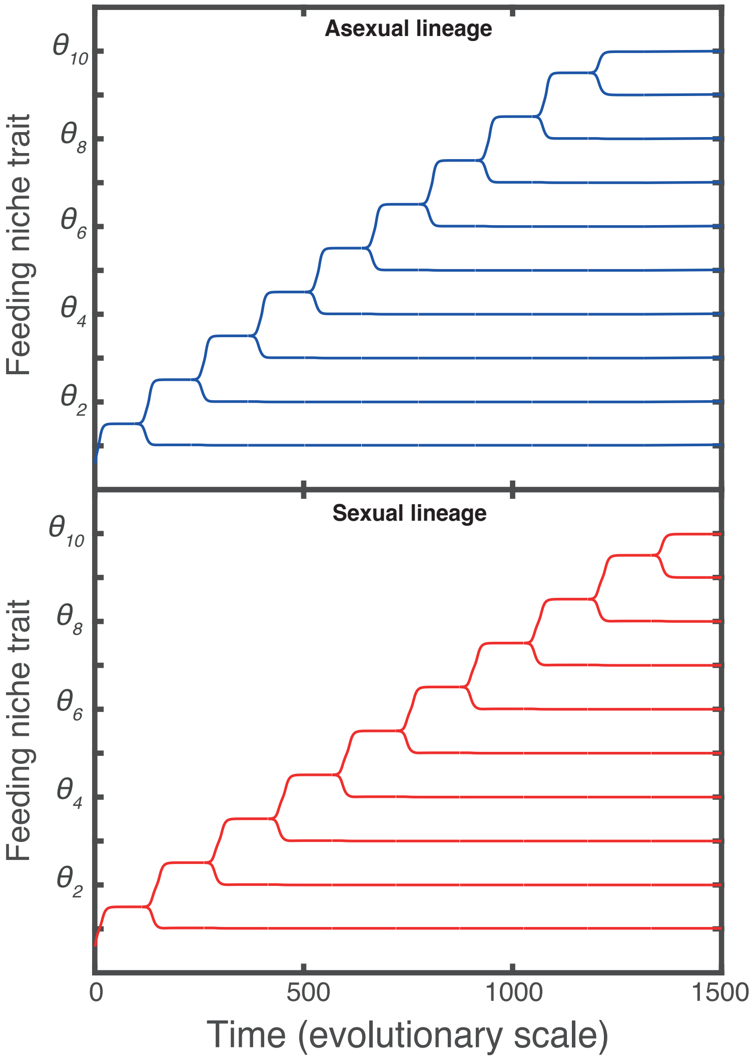

Figure S1. Evolutionary dynamics of an asexual (top) and a sexual (bottom) lineage when only natural selection drives the evolution of the feeding niche trait. Directional natural selection is given by eq. 6 and eq. 8 in the sexual and in the asexual lineage, respectively (see Methods). The ancestral population encouters 10 food resources (optimal trait values to feed on each resource are indicated by the ticks on the vertical axes). Parameter values as in table S1.

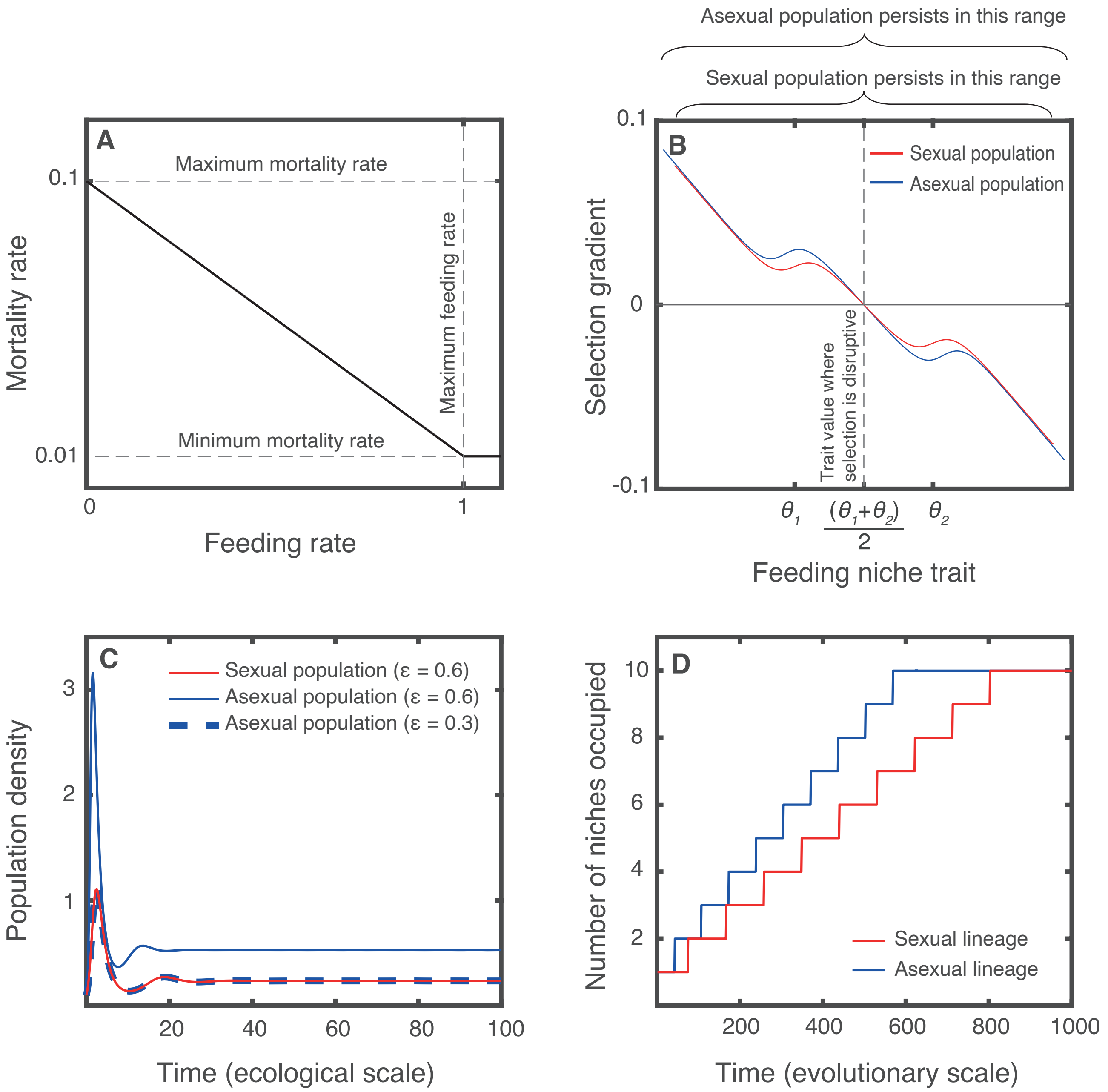

Figure S2. *The effect of natural selection alone on diversification when mortality depends on food consumption*

A) We test the robustness of our results by implementing a mortality function that depends on the rate of food intake. B) Natural selection is stronger in asexual than in sexual populations, in other words, the magnitude of the selection gradient is always larger in an asexual than in a sexual population experiencing the same ecological conditions. C) This is due to the difference in population density: a sexual population (red line) reaches a lower density than an asexual population experiencing the same ecological conditions (blue solid line), and the same density than an asexual population with a half of the efficiency to convert food into offspring (blue dashed line). D) As a result, in a habitat with multiple niches, an asexual lineage diversifies to occupy all niches faster than a sexual lineage. In B and C, a population encounters two food resources (optimal traits to feed on the resources: $\theta_1$= 1 and $\theta_2$= 2). In C, the ecological dynamics are calculated assuming a (nonevolving) trait value of 1.4, and using eq. SI1.2 and eq. 3 for the sexual population and eq. SI1.3 and eq. 4 for the asexual population. In D, the ancestral population encounters ten food resources. In all panels, the carrying capacity of each food resource is equal to 5 g/L. Other parameter values as in table S1.

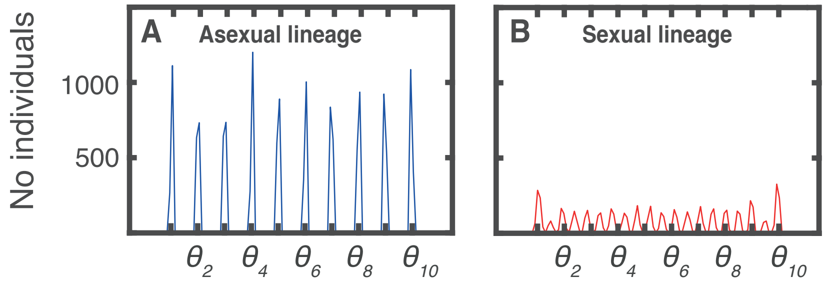

Figure S3. Final trait distribution of the simulation in A) figure 3A and B) 3B.

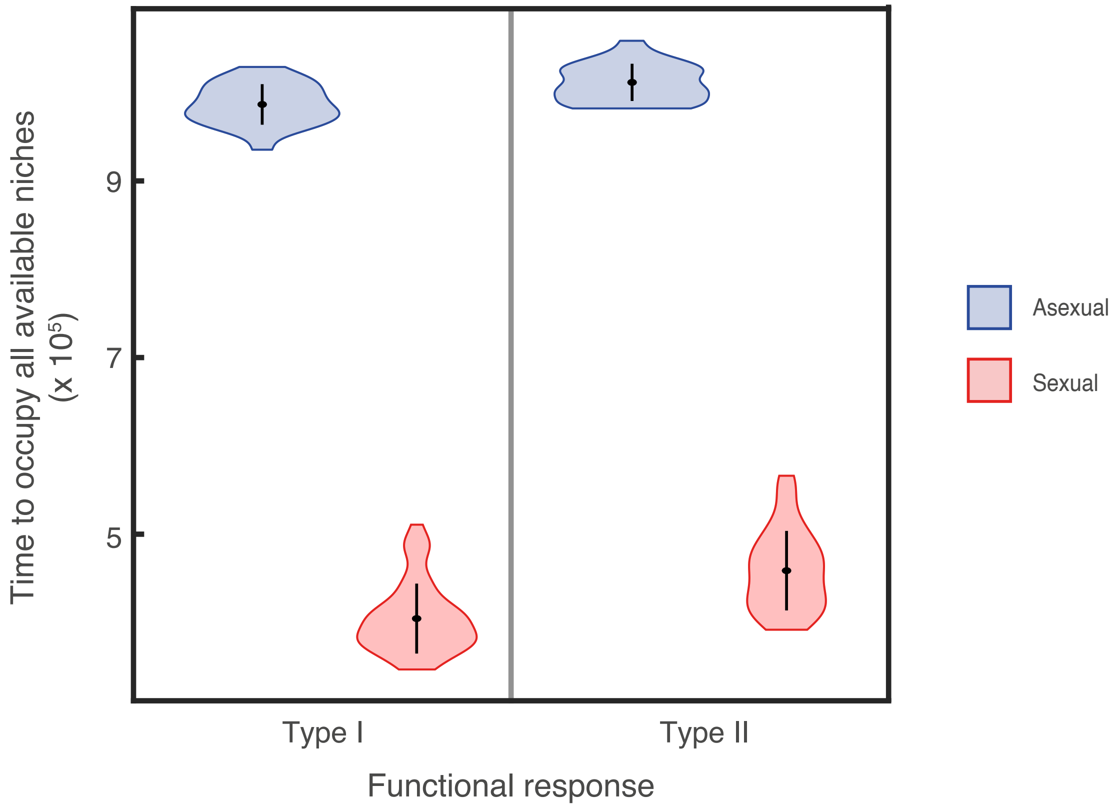

Figure S4. Time required to occupy all available niches for sexual and asexual lineages whose individuals feed following a Type I (left) and Type II (right) functional response. The expansion over the trait space occurs faster in a sexual lineage than in an asexual lineage regardless of the functional response. Mean (black dots) and SD (black lines) in each violin plot correspond to 50 replicate simulations. In the case of functional response Type II, the individual food intake follows eq. SI1.1 (see SI1), with handling time $h_T = 1$. Time in the vertical axis correspond to the number of time steps in the IBM. Other parameter values as in table S1 and S2.

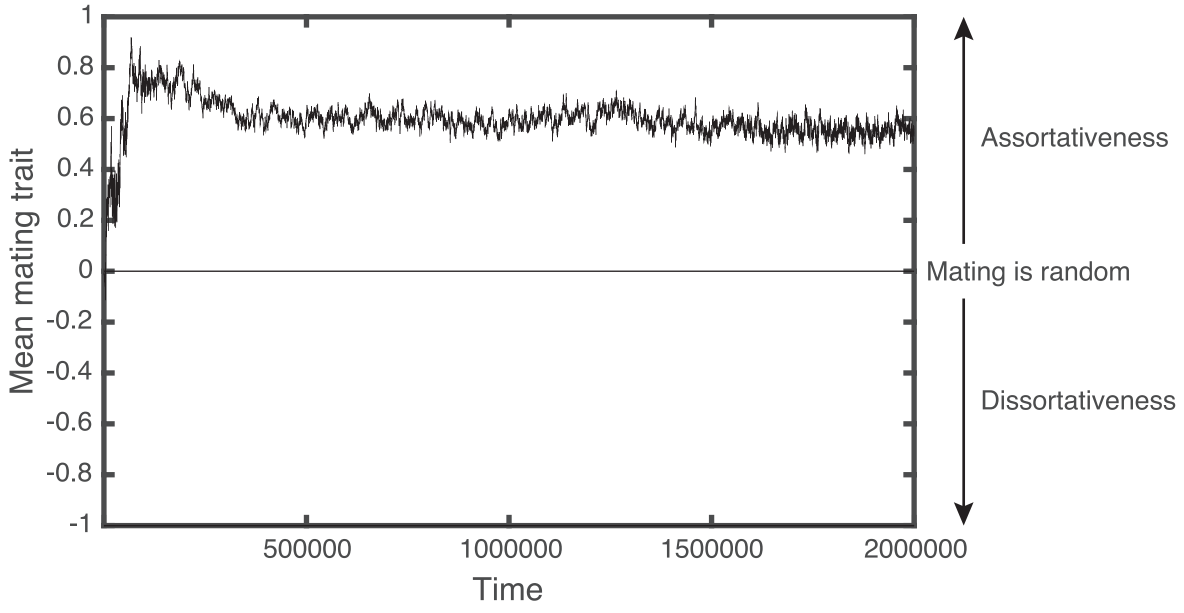

Figure S5. Mating trait dynamics of the simulation in figure 3B. At time t = 0, all individuals have a mating trait equal to 0. Assortativeness is quickly selected for.

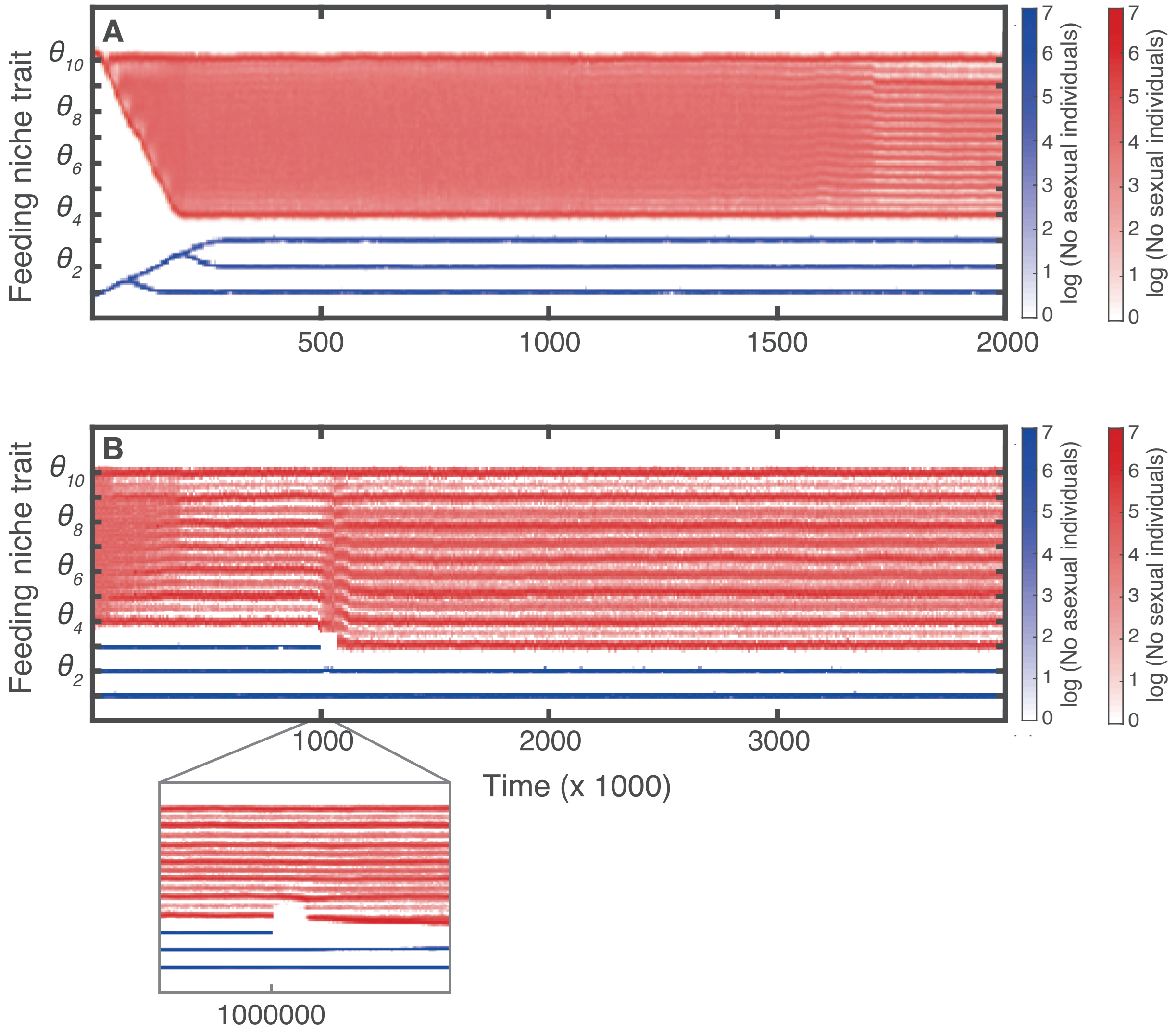

Figure S6. A) Both a sexual (red) and an asexual (blue) populations colonize an environment with 10 available niches. Due to the slower trait expansion of the asexual lineage, this lineage occupies only 3 niches, whereas the sexual lineage occupies 7 niches. B) Simulations of a disturbance event (at time 1000000) that causes the extinction of all individuals with trait value larger than 2.5 and smaller than 4.5 (individuals with trait values near the optimal to feed on resource 3 and resource 4 die) when both the sexual and asexual lineage coexist (see SI7). Ticks in the vertical axes show the trait values corresponding to the optima to feed on food resources. In B, the trait distribution of the lineage at time 0 is the trait distribution at the end of A. Parameter values as in table S1 and S2.

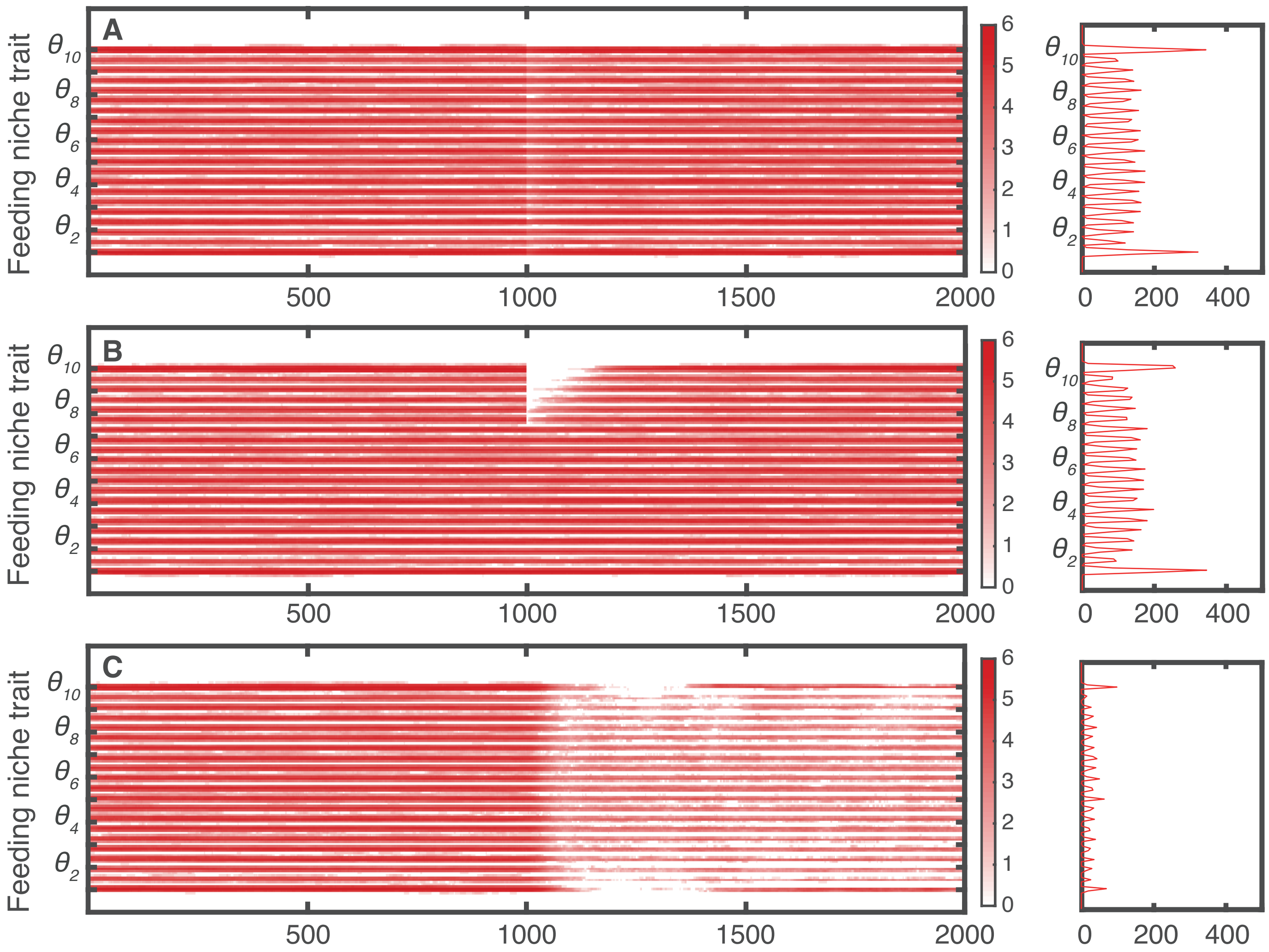

Figure S7. Simulations of diverse disturbance events. After discrete clusters have emerged, we simulate A) an event (time 1000) that kills 50% of individuals randomly, B) an event (time 1000) that causes the extinction of all individuals with trait value larger than 7.5, C) a gradual increase in mortality up to 0.08 per unit of time (four times the original mortality rate) in steps of 0.001 per unit of time (starting at time 1000), and D) an event (time 1000000) that causes the extinction of all individuals with trait value larger than 2.5 and smaller than 4.5 (individuals with trait values near the optimal to feed on resource 3 and resource 4 die) when both the sexual and asexual lineage coexist (see SI7). In A, B and C, the trait distribution of the lineage at time 0 is the trait distribution of the sexual lineage at the end of the simulation shown in figure 3B. Parameter values as in table S1 and S2.

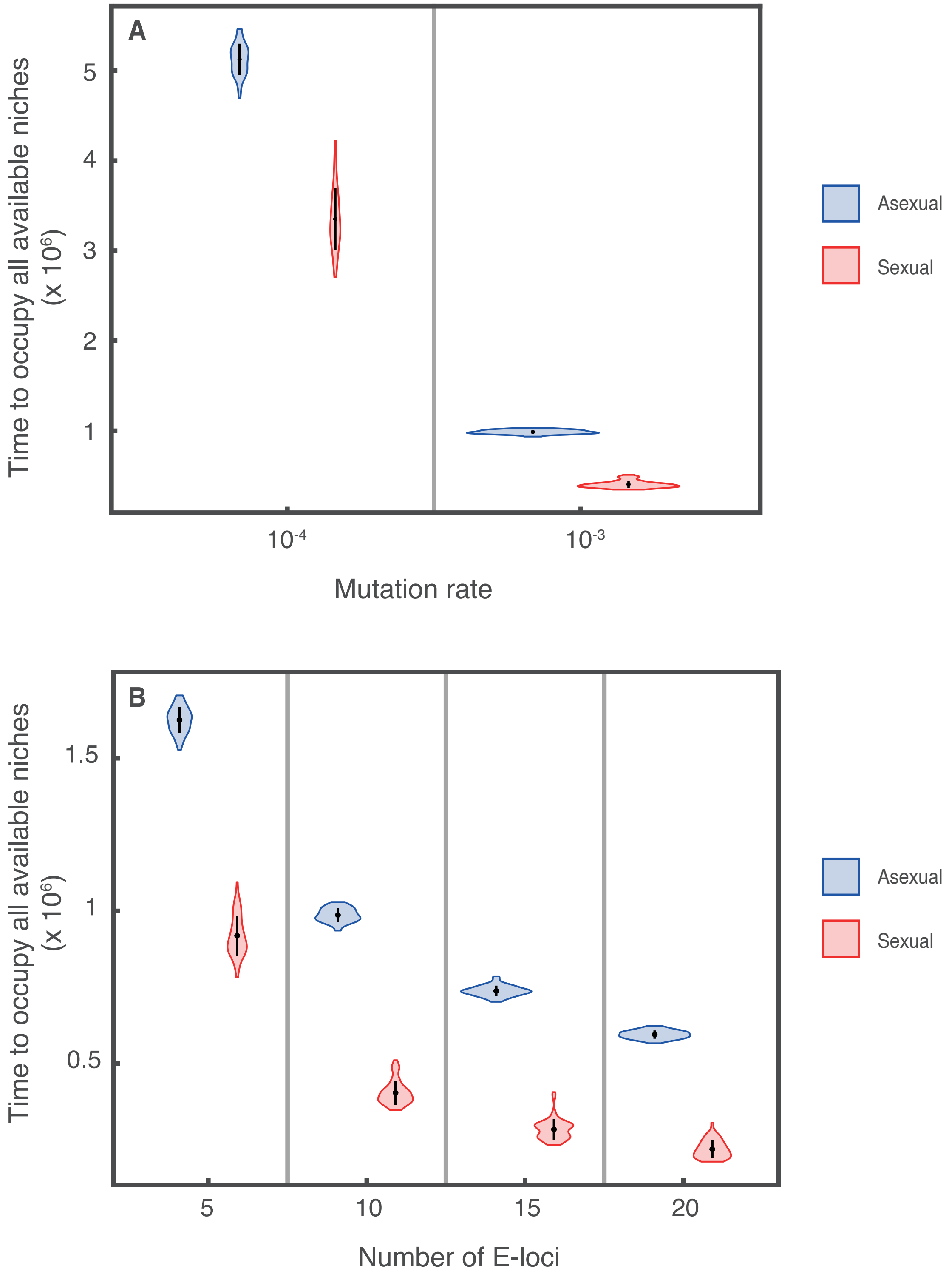

Figure S8. Time required to occupy all available niches for sexual and asexual lineages with different A) mutation rates and B) number of loci encoding the feeding niche trait (E-loci). The expansion over the trait space occurs faster in a sexual lineage than in an asexual lineage under different mutation rates and different number of loci encoding the feeding niche trait. Mean (black dots) and SD (black lines) in each violin plot correspond to 50 replicate simulations. Parameter values as in table S1 and S2.

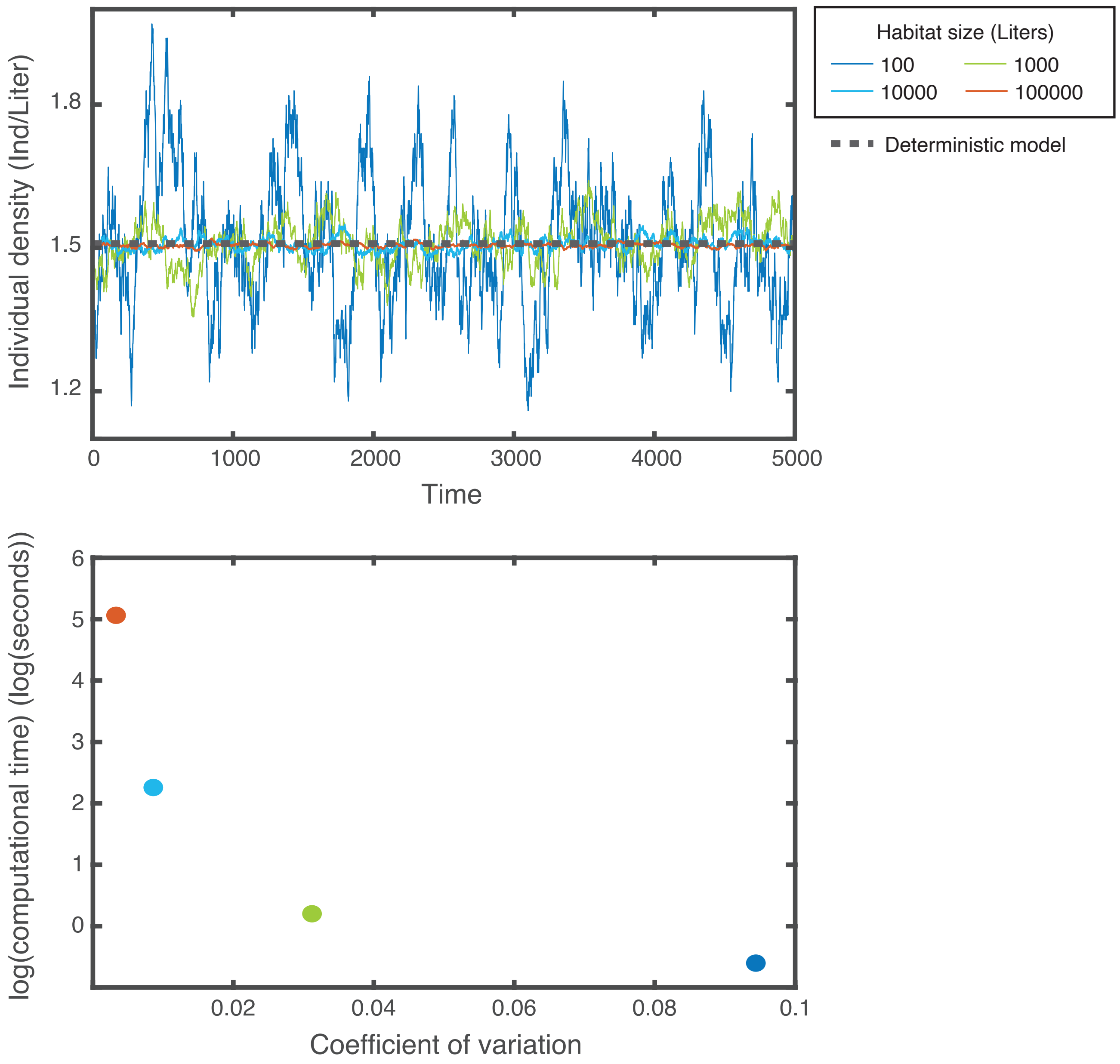

Figure S9. Simulations of the IBM of the asexual lineage for different habitat sizes. While increasing habitat size reduces demographic stochasticity, it also increases the computational time exponentially (notice the logarithmic vertical axis in the bottom plot). The largest reduction in the coefficient of variation (as a proxy for demographic stochasticity) with the smallest increase in computtional time, occurs when the habitat size increases from 100 to 1000. We therefore select 1000 as habitat size for IBM simulations. The four populations simulated reproduce asexually and evolution cannot take place ($\mu_E = 0$). Simulations were initialized using the individual and food resource densities equal to the densities of the equilibrium calculated with the deterministic model (eq SI4.9 - SI4.11 in SI4). Individuals are identical and their feeding niche trait is equal to 0.8. The habitat has 2 food resources ($\theta_1 = 1$, $\theta_2 = 2$) and the productivity is $P = 10$. This productivity level implies that each food resource has a carrying capacity $R_{1\,max} = R_{2\,max} = 5$, which is equivalent to the carrying capacity of each resource in figures of the main text. Other parameter values as in table S1 and S2.

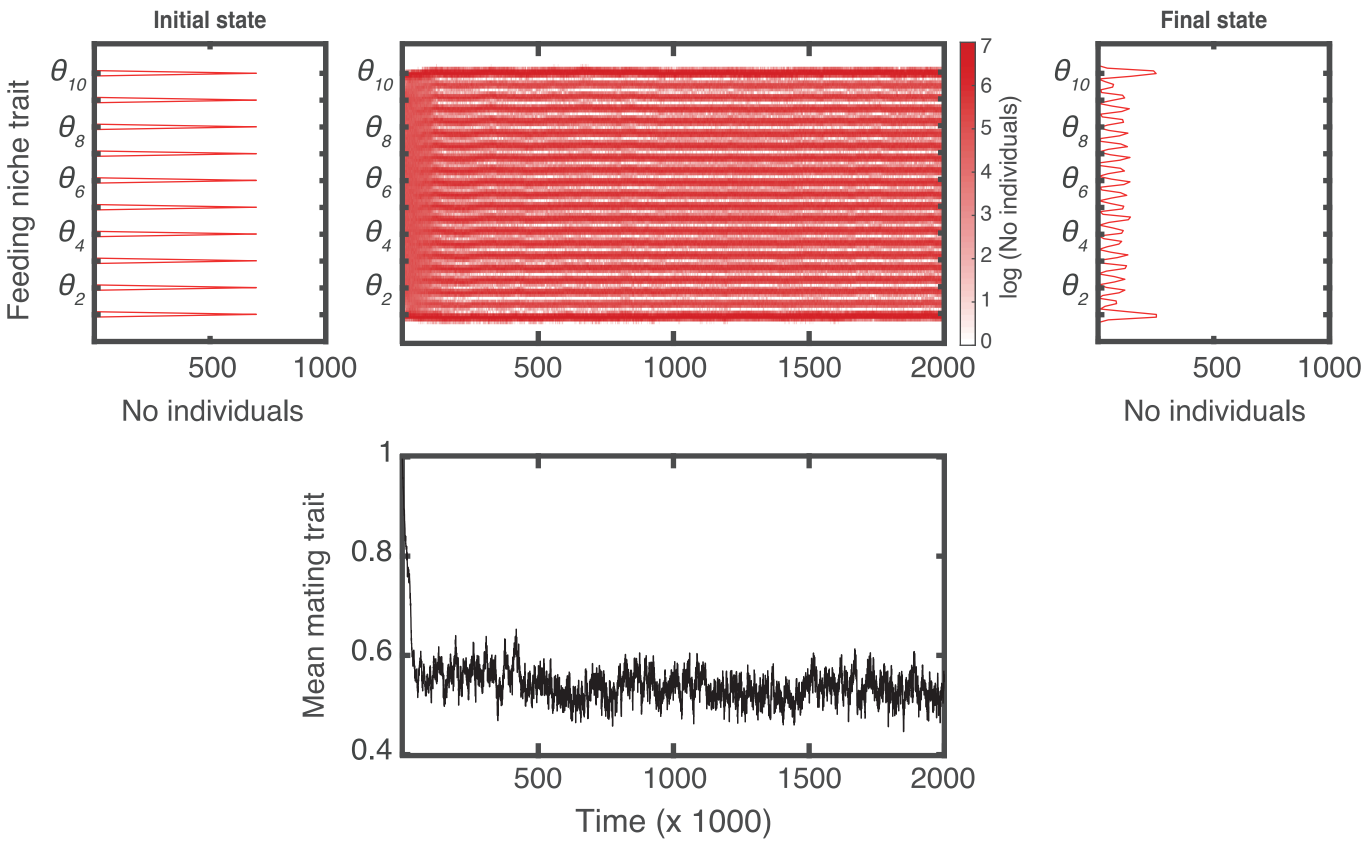

Figure S10. A simulation is initiated with ten populations with feeding niche trait values as shown in the top left panel and mating trait equal to 1 (assortative mating is maximum). These populations quickly collapse into a hybrid swarm and later diverge into discrete generalists and specialists populations. Parameter values as in table S1 and S2.
